# Loss of olfaction reduces caterpillar performance and increases susceptibility to a natural enemy

**DOI:** 10.1101/2024.12.17.629055

**Authors:** Qi Wang, Yufei Jia, Hans M. Smid, Berhane T. Weldegergis, Liana O. Greenberg, Maarten Jongsma, Marcel Dicke, Alexander Haverkamp

## Abstract

Insect herbivores such as caterpillars, are under strong selection pressure from natural enemies, especially parasitoid wasps. Although the role of olfaction in host-plant seeking has been investigated in great detail in parasitoids and adult lepidopterans, the caterpillar olfactory system and its significance in tri-trophic interactions remain poorly understood. In this study, we investigated the olfactory system of *Pieris brassicae* caterpillars and the importance of olfactory information in the interactions among this herbivore, its host plant *Brassica oleracea* and its primary natural enemy *Cotesia glomerata*. To examine the role of olfaction, we utilized CRISPR/Cas9 to knockout the odorant receptor co-receptor (*Orco*). This knockout (KO) impaired olfactory detection and primary processing in the brain. *Orco* KO caterpillars exhibited reduced weight and lost preference for their optimal food plants. Interestingly, the KO caterpillars also experienced reduced weight when challenged by the parasitoid *C. glomerata* whose ovipositor had been removed, and the mortality of the KO caterpillars under the attack of unmanipulated parasitoids increased. We then investigated the behavior of *P. brassicae* caterpillars in response to volatiles from plants attacked by conspecific caterpillars and volatiles from plants on which the caterpillars were themselves attacked by *C. glomerata*. After analyzing the volatile compounds involved in these interactions, we concluded that olfactory information enables caterpillars to locate suitable food sources more efficiently as well as to select enemy-free spaces. Our results reveal the crucial role of olfaction in caterpillar feeding and natural-enemy avoidance, highlighting the significance of chemoreceptor genes in shaping ecological interactions.

## Introduction

Chemoreception plays a critical role in the insect life cycle. For instance, insects exploit chemical cues to locate food plants, to find and select mates, to evaluate oviposition sites, and to avoid natural enemies (Dicke & Grostal, 2001). Olfaction, a pivotal component of chemoreception, involves multiple processes in insects and has been extensively investigated in many species (Haverkamp et al., 2018), including its role in the interactions with conspecifics and natural enemies (Kannan et al., 2022). Thus far, the study of olfaction in insects has largely centered upon imagos, the final life stage of insects. However, chemical information likely also plays an essential role in host plant choice and natural-enemy avoidance during the immature life stage, which are especially pronounced in holometabolous insects. Nonetheless, the olfactory system in larval herbivores remains largely unexplored.

Tri-trophic interactions among plants, herbivores and natural enemies have been widely investigated in the last few decades, and chemoreception is considered to play an important role in these systems (Vet & Dicke, 1992; Turlings & Erb, 2018). In particular, abundant research has demonstrated that olfactory information, such as plant volatiles, is crucial in the coevolution of herbivores and natural enemies (Turlings & Erb, 2018; Dicke & Baldwin, 2010; Haverkamp & Smid, 2020). In addition to plant volatiles, chemicals emitted by the insects themselves also play an important role (Venugopal et al., 2020; Ebrahim et al., 2015). Under strong selection pressure from natural enemies, herbivores have developed several integrated avoidance strategies based on the detection of chemical cues. *Drosophila melanogaster* fruit flies, for example, prefer to lay eggs on citrus fruits because the volatiles emitted by citrus are repellent to the endoparasitoid *Leptopilina boulardi*, a natural enemy of fruit fly larvae (Dweck et al., 2013). Similarly, the brown planthopper *Nilaparvata lugens* deposits eggs on rice plants infested by the striped stem borer *Chilo suppressalis*, which is repellent to the egg parasitoid *Anagrus nilaparvatae* (Hu et al., 2020).

Insect olfaction is dominated by two classes of olfactory receptor genes, odorant receptors (ORs) and ionotropic receptors (IRs). ORs detect the majority of volatiles in the insect habitat. In all ORs the odorant receptor coreceptor (Orco) is required to form a functional ion channel in the membrane of the odorant receptor neurons (ORNs) (Sato et al., 2008; Wicher et al., 2008; Jones et al., 2005). Previous research indicates that silencing the *Orco* gene can lead to a severe olfaction deficiency, causing a loss of social behavior, impaired mating and significantly reduced foraging behavior (Trible et al., 2017; Yan et al., 2017; Fandino et al., 2019). Previous reports have also clearly shown that adult insect behavior and neuronal development were greatly affected when *Orco* is knocked out (Trible et al., 2017; Yan et al., 2017). Yet, potential changes in olfaction-deficient larvae remain largely unexplored and little is known about how olfaction contributes to the co-adaptation of larvae and their natural enemies, especially in the larvae of non-model insects such as lepidopteran caterpillars.

*Pieris brassicae* is a very common butterfly species that causes substantial agricultural production loss across Europe and the Indian sub-continent (Feltwell, 1982; Hasan & Ansari, 2011; Hasan & Ansari, 2010). The butterflies utilize chemoreception to evaluate oviposition sites and host plants (van Loon et al., 1992a; van Loon et al., 1992b; Wang et al., 2023). The caterpillars also have a well-developed olfactory system (Wang et al., 2024) and chemical communication has been demonstrated to be involved in the interaction between herbivores and their natural enemies (Stam et al., 2014; Dicke & Baldwin, 2010). Natural enemies such as parasitoid wasps are highly attracted to volatiles from cabbage plants infested by *P. brassicae* caterpillars (Mattiacci et al., 1994). *P. brassicae* caterpillars are thus under strong selection pressure from natural enemies, though caterpillars can defend themselves by spitting at natural enemies while being attacked (Müller et al., 2003).

To gain comprehensive insight into the role of olfaction in *P. brassicae*, and to understand the role of olfaction in their coevolution with natural enemies, we employed a well-established research system utilizing *Brassica oleracea* as host plants, *P. brassicae* caterpillars as herbivores and *C. glomerata* parasitoid wasps as natural enemies. In this study, we knocked out *Orco* by CRISPR/Cas9 in *P. brassicae* to investigate how olfaction guides the host plant choice and natural-enemy avoidance of this insect herbivore. The knockout was verified in caterpillars by staining Orco and glomeruli in larval antennae and the larval antennal center (LAC) respectively and was further confirmed in butterflies by electrophysiological investigations. We also examined the egg-laying behavior of KO butterflies, which is the basis of establishing a homozygous KO insect colony for our subsequent studies. The development of caterpillars on cabbage plants was evaluated, and the performance of caterpillars under the threat of natural enemies was studied to investigate the ecological significance of larval olfaction in interacting with their natural enemies. The larval host-plant seeking behaviors were further evaluated to determine the importance of olfaction in locating food sources. Volatiles emitted by caterpillars and parasitoids were analyzed to identify candidate chemicals that might play a role in this interaction. Our study provides further knowledge on the olfactory system in insect herbivores and might help to clarify the ecological significance of olfaction in the interaction with their natural enemies.

## Results

### *Orco* knockout and neuronal deficiency

To test for potential off-target regions of our sgRNAs, we searched these sequences against the *Pieris brassicae* genome to identify any such regions. Neither did we find any off-target sites with a less-than-5nt alignment mismatch from Exonerate searching results, nor did we find any off-target sites with a less-than-4-nt alignment mismatch from CHOPCHOP searching results. Furthermore we did not find any developmental differences between wildtype (WT) animals and knockouts (KO) when caterpillars were raised on leaf discs in Petri dishes (Fig S1) or in the hatching rate of fertilized WT and KO eggs (Fig S2). Taken together these results indicate that no off-target sites were available for our sgRNA. Through screening all butterflies that were reared from injected eggs using PCR and sequencing, we identified an *Orco* mutant line in *P. brassicae*. This mutant line exhibited an 11-bp insertion and a four-bp deletion in the second exon, resulting in a shifted reading frame (Fig 1A). The Orco amino acid sequence showed a significant change mediated by an early stop codon (Fig 1B). These results indicate that the KO of the *Orco* gene was successful.

**Fig 1.**
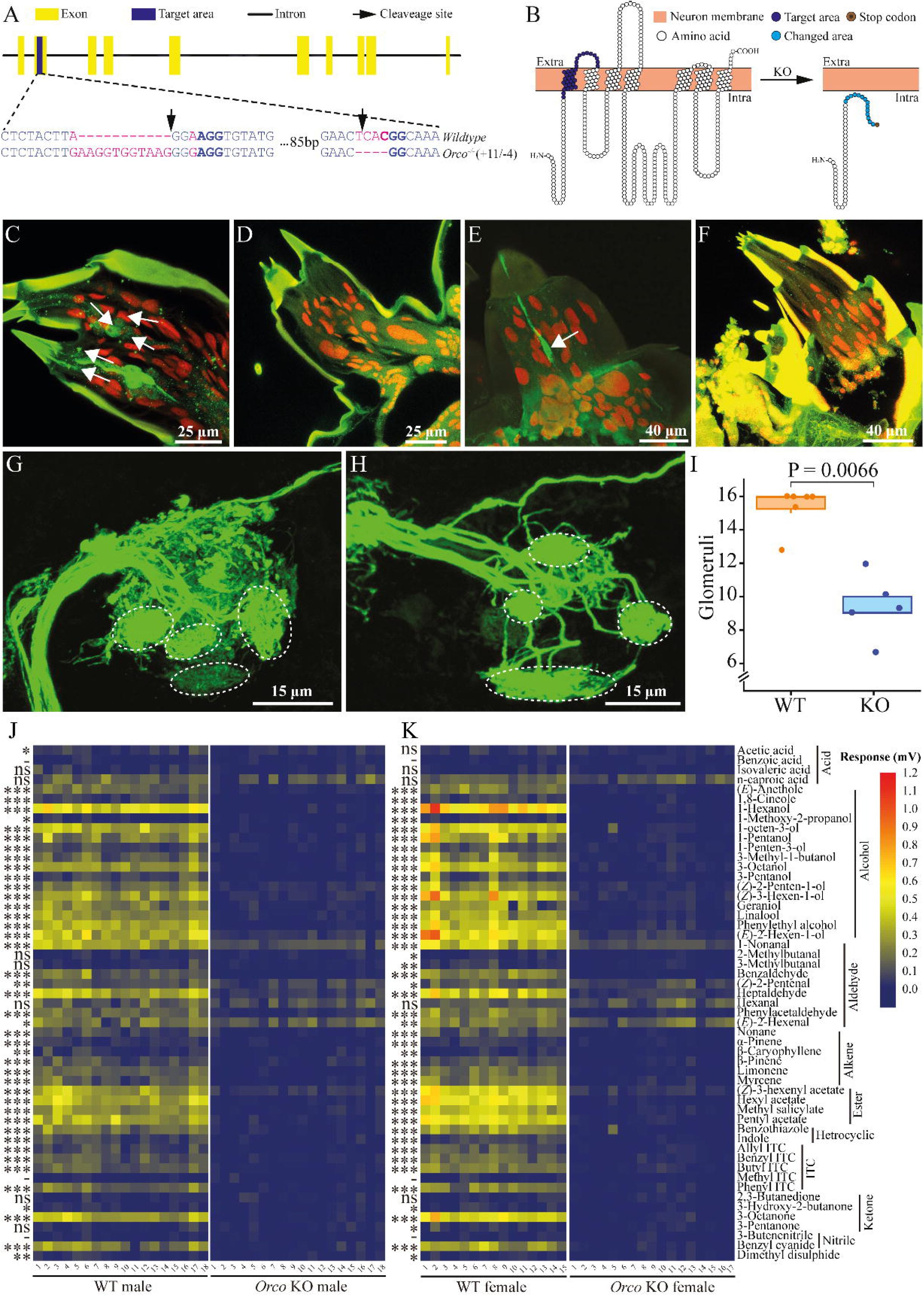
*Orco* knockout by CRISPR/Cas9 and verification in *Pieris brassicae*. (A), gene structure of *Orco*. Yellow blocks indicate exons and blue block indicates target area in the second exon. Black solid line indicates intron. Black arrows indicate the designated cleavage site of the Cas9 protein. The target knockout area is magnified to show the sequence. Blue letters indicate base pairs in the second exon segment, purple letters indicate mutation sites. Protospacer adjacent motif (PAM) sequences are in blue bold. (B), predicted transmembrane structure of Orco. The left and right panels represent wildtype (WT) Orco and mutated Orco transmembrane domains, respectively. Orange blocks indicate ORN membrane, extracellular (Extra) and intracellular (Intra) are shown. White circles indicate amino acids of Orco, blue circles on the left panel indicate the target mutation area. Cyan circles and the brown circle on the right panel indicate the mutated area and the early stop codon, respectively. Orco-positive ORNs and larval antennal center. *P. brassicae* larval antennae and larval antennal center (LAC) staining. (C), Odorant receptor neurons (ORNs) were stained (green cells, several are indicated by white arrows) in WT larval antennae. (D), ORNs were not stained in *Orco* knockout (KO) larval antennae. (E), An ORN was stained (green cell, indicated by white arrow) in WT larval palps. (F), ORNs were not stained in KO larval palps. (G), glomeruli in the LAC of a WT brain. (H), glomeruli in the LAC of a KO brain. Glomeruli are indicated by white dashed circles in (G). In (H), the white dashed lines indicate the approximate position of missing glomeruli (not directly corresponding to panel (G)). (I), number of glomeruli counted in WT caterpillar brain (orange) and *Orco* KO caterpillar (blue) brain. A significant difference was detected by Wilcoxon rank-sum test. (J), electroantennogram (EAG) response of male butterflies (n = 18 for both WT and KO). (K), EAG response of female butterflies (n = 15 for WT and n = 17 for KO). Left panels represent wildtype (WT) butterfly EAG responses, and right panels represent are knockout (KO) butterfly EAG responses. Significant differences between WT and KO butterflies were identified using Student’s t-test when the data was normally distributed and the variances were equal or using a Kruskal-Wallis rank-sum test when these criteria did not apply. Significance levels are indicated by asterisks, ns (P > 0.05), * (0.01 < P < 0.05), ** (0.001 < P < 0.01), *** (P < 0.001), - (no response recorded from WT butterfly antennae). Butterfly antennal responses (mV) to the tested chemicals are indicated by a color scale from navy (0 mV) via yellow to red (1.2 mV).

To further validate the KO in *P. brassicae* and investigate the neuronal mechanism underlying olfactory deficiency in KO caterpillars, we performed staining of Orco in third-instar larvae (L3) using a lepidopteran specific Orco antibody, which has been validated previously (Fandino et al., 2019; Wang et al., 2024; Nolte et al., 2016). In L3 WT caterpillar antennae, a total of 34 ORNs were successfully identified (Wang et al., 2024) (Fig 1C). However, no Orco-positive ORNs were found in the antennae of KO caterpillars (Fig 1D). Similarly, we detected a low number of Orco-positive neurons in the maxillary palps of L3 WT caterpillars (Fig 1E), but no Orco-positive signal was detected in the maxillary palps of KO caterpillars (Fig 1F). By tracing the ORN axons to their glomeruli in the brain, we observed some differences between WT and *Orco* KO caterpillars. In the larval antennal center (LAC), glomeruli displayed well-defined shapes with clear boundaries in the WT LAC (Fig 1G). However, in the KO LAC, the glomeruli were less clearly separated (Fig 1H). Only 7-12 glomeruli were identified in KO LAC, which was a significant reduction (P = 0.0066, Wilcoxon rank-sum test) compared to WT LAC (Fig 1I).

### Butterfly electrophysiological response to chemical compounds and oviposition

We then conducted additional tests comparing the antennal electrophysiological responses of male and female butterflies from both genotypes using electroantennography (EAG), to further confirm the KO of *Orco* in butterflies. *Orco* KO butterflies of both sexes displayed a complete loss of responses to the tested chemicals with the exception of isovaleric acid, n-caproic acid, hexanal and 2,3-butanedione, where no significant difference was detected between KO and WT butterflies. Reduced responses were recorded in KO butterflies to 1-nonanal, (*Z*)-2-pentenal, (*E*)-2-hexenal and (*Z*)-3-hexenyl acetate. Besides, WT male (n = 18) and female (n = 15) butterflies responded similarly to most of the volatile compounds with a few exceptions where WT females showed significantly different responses to 2-methylbutanal (P = 0.0183, Kruskal-Wallis test), 3-methylbutanal (P = 0.0030, Kruskal-Wallis test) and 3-pentanone (P = 0.0263, Student’s t-test) compared to KO females (n = 17), and WT males (n = 18) responded significantly stronger to acetic acid than KO males (P = 0.0217, Student’s t-test). Moreover, both WT males and females showed no response to benzoic acid, methyl ITC and 3-butenenitrile (P > 0.05, Student’s t-test) (Fig 1J and 1K).

To investigate the role of olfaction in butterfly mating and oviposition, and evaluate the feasibility of establishing homozygous *Orco* KO colony for subsequent larval studies, pairs of newly emerged butterflies were collected and reared in a cage and allowed to mate freely. Upon dissection, it was found that KO butterflies (n = 14) mated fewer times than WT butterflies (P = 0.0199, n = 11; Wilcoxon rank-sum test), since the majority of the KO butterflies (9 out of 14) did not mate during the experiment (Fig S3). Out of the five KO butterflies that did mate, all but one mated singly. Of the WT butterflies, eight out of 11 mated, with most mating twice and one individual even thrice. Female KO butterflies also deposited fewer eggs than WT butterflies during the experiment (P < 0.0001, GLM negative binominal) (Fig S4). To determine the hatching rate, the number of newly hatched caterpillars of both genotypes were counted. It was found that virgin butterflies of both genotypes did not deposit any fertilized eggs, while mated butterflies exhibited similar fecundity with comparable hatching rates between the two genotypes without any significant difference (Fig S2).

### Caterpillar performance on cabbage plants

To assess the ecological significance of knocking out *Orco* in *P. brassicae* caterpillars, we first conducted a comparison of caterpillar growth between the two genotypes on their optimal host plant. The results show that in the absence of parasitoids, WT caterpillars exhibited better performance, gaining more weight compared to KO caterpillars (P = 0.0268, n = 17 for both genotypes; Student’s t-test) (Fig 2A). However, we did not find significant survival difference between the two genotypes while collecting caterpillars after the experiment (WT mortality = 8.8 %, n = 17; KO mortality = 2. 9 %; P = 0.0880, n = 17; Wilcoxon test). Nor was there a developmental difference when caterpillars were reared in a Petri dish supplied with leaf discs (P=0.97, n = 10 for both genotypes; Wilcoxon test) (Fig S1). Subsequently, to determine if the differences in feeding and survival are dependent on food-plant location, a two-choice assay was conducted with cabbage leaf discs, tomato leaf discs and green paper discs in a glass Petri dish to investigate the role of olfaction in host-plant selection (Fig 2B). Both genotypes of caterpillars chose the cabbage leaf disc significantly more often than the green paper discs (P = 0.0016 for KO, χ^2^ = 9.92; P < 0.0001 for WT, χ^2^ = 21.41; Chi-square test), and no difference was detected between the two genotypes (P = 0.1460, χ^2^ = 2.11; Chi-square test). Similarly, when the cabbage leaf disc was replaced with a tomato leaf disc, both genotypes preferred the tomato leaf disc over the paper disc (P = 0.0143 for KO, χ^2^ = 6.00; P = 0.0009 for WT, χ^2^ = 10.97; Chi-square test) and no difference was found between the two genotypes (P = 0.5477, χ^2^ = 0.36; Chi-square test). However, when caterpillars were provided with both a cabbage leaf disc and a tomato leaf disc, a contrasting preference was observed between the two genotypes (P = 0.0004, χ^2^ = 12.68; Chi-square test). KO caterpillars significantly preferred the tomato leaf disc (P = 0.0466, χ^2^ = 3.96; Chi-square test), while WT caterpillars preferred the cabbage leaf disc (P = 0.0028, χ^2^ = 8.97; Chi-square test).

**Fig 2.**
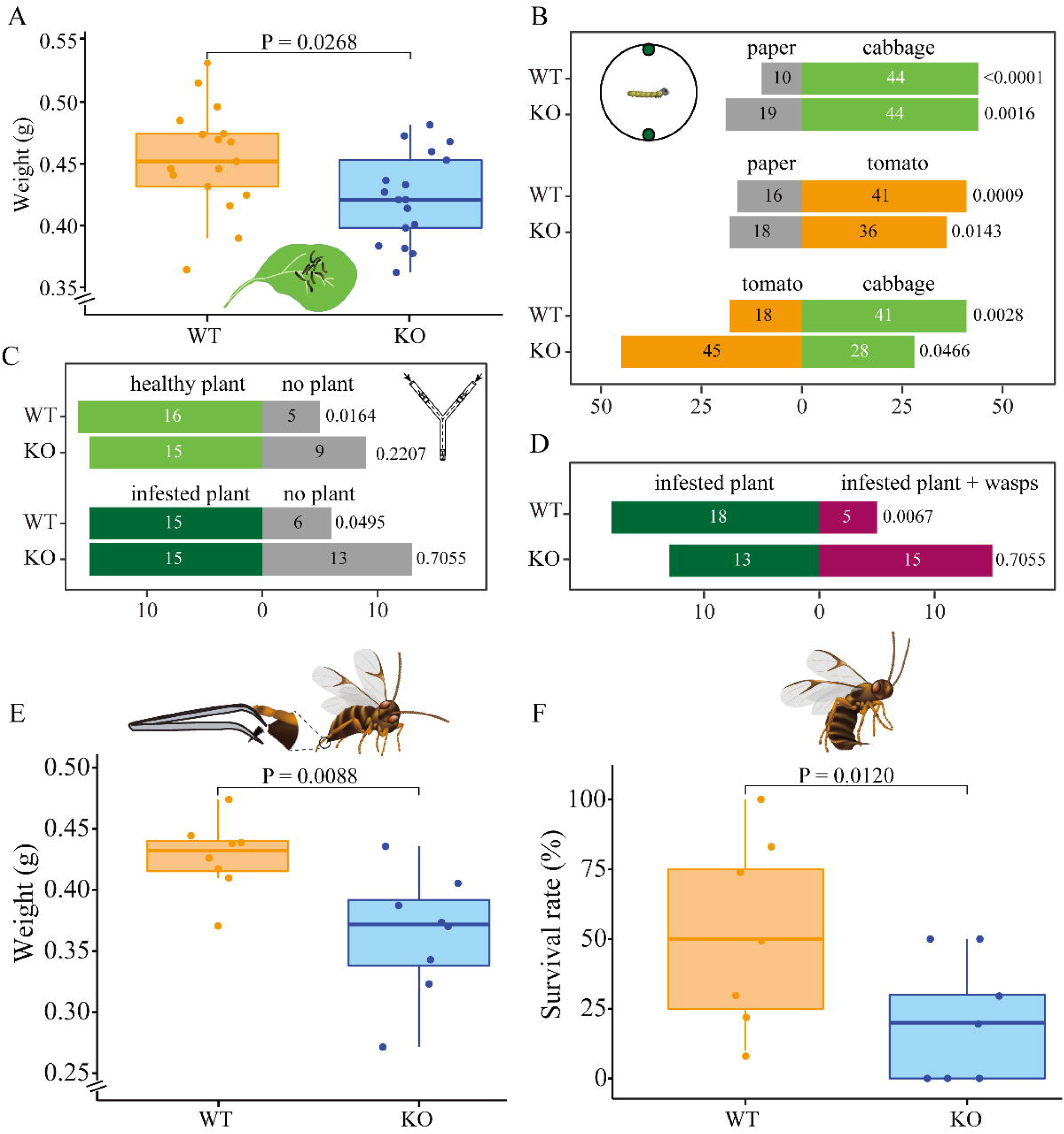
*Pieris brassicae* caterpillar growth and foraging behavior. (A), *P. brassicae* caterpillar growth on cabbage plants, y axis shows weight (g) of WT and KO caterpillars after 10 days of feeding , n = 17. (B), caterpillar behavioral choices in a Petri dish. The numbers of caterpillars that chose one of the two discs are shown in the respective bars (n = 54 -73). Schematic drawing shows behavioral setup. Petri dish diameter was 188 mm, disc diameters were 13 mm. Discs in the Petri dish represent cabbage, paper and tomato leaf discs. Grey bars indicate paper disc choices, light green indicates cabbage leaf disc choices and orange indicates choices for tomato leaf discs. (C), caterpillar behavioral choices in Y-tube olfactometer without parasitoid wasps. Schematic drawing shows the Y-tube olfactometer. The dashed line indicates a black metal Y wire in the center of glass Y-tube olfactometer. The main arm of the Y-olfactometer is 200 mm length, the lateral arms are 275 mm length, the angle between the lateral arms is 80 °. Light green, dark green and grey bars represent choices for healthy plant, infested plant and no plant respectively. Healthy plant, plants were not treated; no plant, an empty jar without any insect or plant; infested plant, plants were infested by early L3 caterpillars. (D), caterpillar behavioral choices in Y-tube olfactometer with parasitoid wasps. Different treatments are in different colors. Dark green, infested plant - wasps; magenta, infested plant + wasps. Significant differences were tested between wildtype (WT) and knockout (KO) caterpillars or between two discs by Chi-square test in panel B-D, P-values are shown on the right side of each bar. (E), *P. brassicae* caterpillar growth when exposed to disarmed *C. glomerata* female parasitoids, y axis shows weight (g) of caterpillars after 10 days of feeding, n = 8. Schematic drawing shows disarmed female *C. glomerata* (ovipositor removed). (F), *P. brassicae* caterpillar survival rate when exposed to healthy *C. glomerata* female wasps, n = 7. Significant differences in development and survival rate were assessed using a one-tailed Student’s t-test, P values are indicated above boxplots. Schematic drawing shows a healthy female *C. glomerata* (unmanipulated). In both panels orange boxplots indicate WT caterpillars, blue boxplots indicate *Orco* KO caterpillars.

We then aimed to determine whether caterpillars would orient towards healthy plants, plants infested with conspecifics and plants on which conspecifics are attacked by parasitoid wasps, using a Y-tube olfactometer. We first compared the behaviors of both WT and *Orco* KO caterpillars in response to clean air, volatiles from a healthy plant and volatiles from a caterpillar-infested plant (Fig 2C). Both WT and *Orco* KO caterpillars tended to prefer a healthy cabbage plant to a clean air control, however WT caterpillars were significantly attracted by volatiles from a healthy plant (P = 0.0164, χ^2^ = 5.76; Chi-square test) while the preference for *Orco* KO caterpillars was not significant (P = 0.2207, χ^2^ = 1.50; Chi-square test). Subsequently, we challenged the caterpillars with volatiles from a infested plant and a clean air control and found that WT caterpillars prefer volatiles from the infested plant to the control (P = 0.0495, χ^2^ = 3.86; Chi-square test), while *Orco* KO caterpillars did not show any preference between the two provided options (P = 0.7055, χ^2^ = 0.14; Chi-square test). We further compared larval host-plant seeking behaviors with the presence of natural enemies on caterpillar-infested plants (Fig 2D). WT caterpillars showed a significant preference for infested plants without parasitoid wasps *C. glomerata* (P = 0.0067, χ^2^ = 7.35; Chi-square test), while *Orco* KO caterpillars did not show any preference for either odor source (P = 0.7055, χ^2^ = 0.14; Chi-square test).

To understand the ecological significance of the odor-guided behavior for the natural enemy avoidance in *P. brassicae* caterpillars, we then evaluated caterpillar performance under the potential threat of natural enemies by exposing the caterpillar to parasitoids after removing the ovipositor of the female wasps (Fig 2E). The results show a significantly higher weight in WT caterpillars than in KO caterpillars (P = 0.0088, n = 8 for both genotypes; Student’s t-test). To further explore the effect of knocking out *Orco* on the interaction of the caterpillars with their natural enemies, we compared the survival rates of caterpillars when exposed to two mated unmanipulated female *C. glomerata* wasps. The findings show that KO caterpillars had significantly lower survival rates compared to WT caterpillars (P = 0.0120, n = 7 for both genotypes; GLM beta-binomial) (Fig 2F).

### Volatile headspace composition

Furthermore, we investigated if caterpillar- and parasitoid-derived chemicals are involved in the behavioral responses. We identified a total of 45 chemical compounds among the five treatments belonging to seven classes, including alcohols, aldehydes, aromatics, ketones, nitrogen and/or sulfur-containing chemicals, terpenoids and others (Table S3). The contribution of identified chemicals towards separating different treatment groups was analyzed using principal component analysis (PCA). This analysis showed a significant separation of most groups with PC 1 carrying 65.4 % of the variance (P = 0.001, PERMANOVA). *Pieris brassicae* caterpillars (Pb, n = 10), *P. brassicae* caterpillar spit (Pb-S, n = 12), *C. glomerata* female wasps (Cg, n = 12) and caterpillar frass (Pb-Fr, n = 10) treatment groups were clearly separated (Pb vs. Cg, Pb vs. Pb-S, Pb vs. Pb-Fr, Pb-S vs. Cg and Pb-Fr vs. Cg, P= 0.0013; Pb-S vs. Pb-Fr, P= 0.0022 pairwise PERMANOVA, Fig 3A), whereas the identified and measured volatile blend from Pb was similar to that of the *P. brassicae* caterpillars-*C. glomerata* parasitoids combination (Pb-Cg, n = 14) (P = 0.0330, pairwise PERMANOVA). The amount of volatiles among the five treatments was also compared. Most of the identified compounds were found in significantly higher amounts in Pb-S and/or Pb-Fr compared to the other treatments (Fig 3B, Table S3). We also found that some chemicals are strongly correlated. The identified chemicals are classified into several subgroups in the hierarchical clustering heatmap according to the abundance among treatments. For instance, the seven chemicals (*Z*)-3-hexen-1-ol, 1-penten-3-ol, dimethyl disulfide, methyl (methylthio) methyl sulfide, dimethyl trisulfide, 2-methylbutanal and 3-methylbutanal formed a clade. Similarly, the six chemicals, 3-methylbutanoic acid, 1-pentanol, benzyl cyanide, 3-(methylthio) propanal, (*Z*)-2-hexen-1-ol and phenylacetaldehyde formed another clade. The clustering analysis demonstrated that chemicals within the same clade are highly correlated. The multivariate data analysis (MVDA) also showed similar clustering when comparing the different treatments (Fig S5-S9).

**Fig 3.**
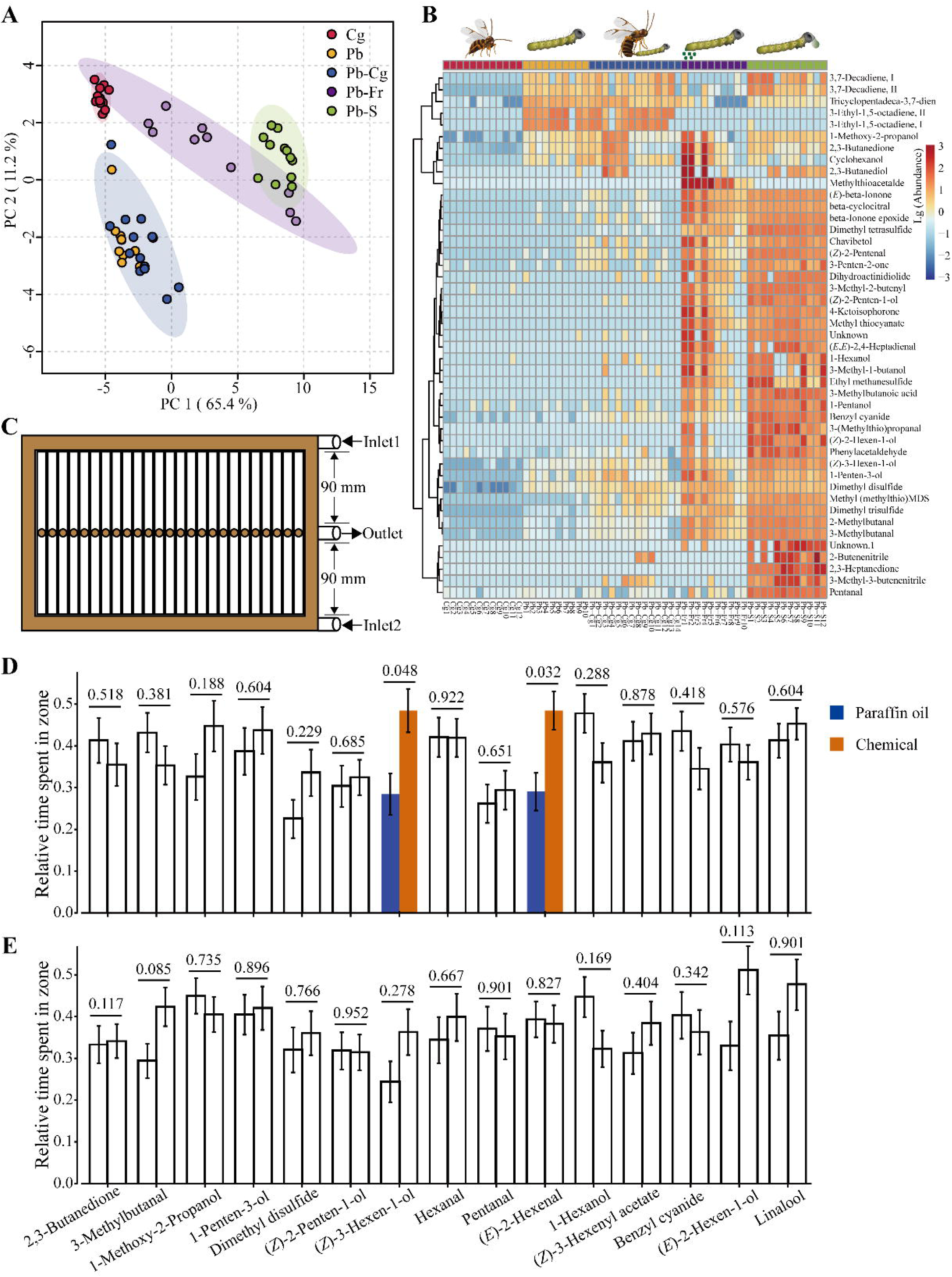
Overview of caterpillar- (*Pieris brassicae*) and parasitoid wasp- (*Cotesia glomerata*) associated volatile compounds. (A), PCA (Principal Component Analysis) two-dimensional score plot of five treatment groups: Cg, *Cotesia glomerata* female parasitoid wasps (n = 12); Pb, *Pieris brassicae* caterpillars (n = 10); Pb-Cg, *P. brassicae* caterpillars in the presence of *C. glomerata* female parasitoid wasps (n = 14); Pb-Fr, *P. brassicae* caterpillar frass (n = 10); and Pb-S, *P. brassicae* spit (n = 12), based on their volatile blend composition. (B), hierarchical clustering heatmap showing the abundance of each identified volatile compound of each treatment. Clustering chemicals in the heatmap indicate higher correlation. (C), schematic drawing of the custom-designed multi-channel arena. (D) wildtype (WT) caterpillar behavioral preference to the tested 15 odorants. (E), *Orco* knockout (KO) caterpillar behavioral preference to the tested 15 chemicals. Unfilled bars represent tests that exhibit no difference between paraffin oil and the odorant of interest. Blue bars indicate the cumulative duration ratio that caterpillars stayed in the paraffin oil zone, dark orange bars indicate the cumulative duration ratio that caterpillars stayed in the odorant zone. Relative time spent in odorant zone = Cumulative duration in a specific zone / Cumulative duration in the arena. Error bars indicate standard errors. In (D) and (E), differences were tested between the two zones using a Wilcoxon rank-sum test, P values are presented above bars (n = 31-50). For each comparison, left bar represents paraffin oil and right bar represents chemical compound.

Based on the results of our chemical analysis, we selected 11 candidate chemicals (1-hexanol, 1-methoxy-2-propanol, 1-penten-3-ol, 2,3-butanedione, 3-methylbutanal, benzyl cyanide, dimethyl disulfide, pentanal, (*E*)-2-hexen-1-ol, (*Z*)-2-penten-1-ol and (*Z*)-3-hexen-1-ol) with high variable importance in the projection (VIP) scores (> 1.0) (Table S3-S8), and another four common plant volatiles (linalool, (*Z*)-3-hexenyl acetate, hexanal and (*E*)-2-hexenal). These odorants were chosen to test the behavioral response of L3 caterpillars. Using a custom-made multiple channel setup (Fig 3C), we conducted behavioral experiments with the caterpillars. The results showed that WT caterpillars displayed a preference for two plant chemicals, namely (*Z*)-3-hexen-1-ol and (*E*)-2-hexenal over paraffin oil (P = 0.0482 and P = 0.0321) (Fig 3D). In contrast, KO caterpillars did not exhibit any significant attraction or avoidance behavior towards the tested chemicals (Fig 3E). Additionally, we found that WT caterpillars were more active in the arena, moving around (Fig S10), while KO caterpillars were less active in the arena, often remaining at the same position within the channels (Fig S11).

## Discussion

Chemical communication plays a significant role in the interaction between plants, herbivores and their natural enemies. Infested host plants emit herbivore-induced plant volatiles (HIPVs) and thereby attract natural enemies such as parasitoid wasps (Mattiacci et al., 1994). While the role of adult herbivores’ olfaction in mediating these tri-trophic interactions has been investigated, the significance of caterpillar olfaction in tri-trophic interactions of plants, herbivores and their natural enemies has remained largely underexplored. In this study, we severely impaired the olfaction of *P. brassicae* by knocking out *Orco* using CRISPR/Cas9 to investigate the ecological importance of olfaction in host-plant choice and enemy avoidance by *Pieris brassicae* larvae. The knockout of olfaction was successful, as we did not find Orco-proteins in the antennae of *Orco* KO caterpillars and showed an altered brain structure in the *Orco* KO caterpillars in comparison to WT caterpillars. Furthermore, the near complete lack of response to odors by *Orco* KO butterfly antennae compared to WT confirms that olfaction was largely lost. Intriguingly, although mating and oviposition were significantly reduced, KO butterflies still produce some homozygous offspring. After this characterization of our mutant line, we aimed to test the role of larval olfaction in a multitrophic framework. We found that the loss of olfaction had a significant effect on the foraging behavior of the caterpillars, which exhibited a reduced growth rate, and a greater vulnerability to the parasitoid wasp *C. glomerata*.

The perception of chemical cues is mediated by the chemosensory signal transduction in insect antennae and further processing of the information in the brain. In *P. brassicae* KO caterpillar antennae, no Orco protein was detected, and chemical signals can therefore not be converted into electrical signals (Sato et al., 2008; Wicher et al., 2008). Interestingly, the non-functional Orco decreased the number of glomeruli in the LAC, indicating that Orco is likely to play a role in the glomerular organization of the LAC (Fig 1C-I). Similar changes have been reported for adult insects of a few hymenopteran species: the ants *Harpegnathos saltator* and *Ooceraea biroi* and the honeybee *Apis mellifera*, where the number and total size of glomeruli decreased while the volume of individual glomeruli increased in *Orco* KO mutants (Trible et al., 2017; Yan et al., 2017; Chen et al., 2021). In contrast, the antennal lobe morphology of *D. melanogaster* was not significantly changed when *Orco* was knocked out (Larsson et al., 2004). In Lepidoptera, the size and number of glomeruli in *Helicoverpa armigera* were found to be comparable between WT and KO adult moths without any significant difference (Fan et al., 2022). However, silencing the pheromone receptor, *SlitOR5* led to a decreased size of a glomerulus in the adult moth *Spodoptera littoralis* and knocking out *Orco* also led to a reduced size of OR-related glomeruli in adult *Manduca sexta* (Koutroumpa et al., 2022; Fandino et al., 2019). The loss of olfactory glomeruli in the caterpillar might, therefore, either indicate that the formation of glomeruli in the embryo is mediated by a different mechanism than during pupation (Oland & Tolbert, 2011), or that the loss of neuronal activity reduces axonal innervations and thereby the size of the glomeruli to an extent that they could no longer be visualized by our method. However, the mechanism of glomerular formation and the roles that olfactory genes play in this process needs to be further investigated in both immature and adult insects (Williams et al., 2022).

The role of olfaction in the behavioral ecology of adult insects has already been studied in quite some detail, and silencing *Orco* has been an important tool for unravelling the importance of chemical communication in adults of various insect species. For instance, in *Orco* KO adults of several insect species, there was a significant reduction in the electrophysiological responses to volatile chemicals such as alcohols and esters (Fandino et al., 2019; Sun et al., 2020). Consequently, *Orco* mutants exhibited notable changes in host-plant location and severely reduced fecundity due to impaired pheromone perception and mating, which also resulted in significantly reduced egg-laying behavior (Fan et al., 2022; Fandino et al., 2019; Liu et al., 2023). Similar to other species, *Orco* KO *P. brassicae* butterflies exhibited a loss of electrophysiological responses to most plant volatiles we tested (Fan et al., 2022; Sun et al., 2020) (Fig 1J and 1K), and the mating behavior of *P. brassicae Orco* KO butterflies was also disrupted (Fig S3). However, we observed some differences: *P. brassicae Orco* KO butterflies were still able to mate to a limited degree (Fig S2). Whereas *Orco* KO adults in some other insect species did not exhibit any mating behavior (Fandino et al., 2019; Fan et al., 2022). This finding suggests that additional sensory systems beyond olfaction, such as vision, may be involved in the mating processes of this day-active butterfly (Carpenter & Sparks, 1982; Obara et al., 2008). Interestingly, while many *P. brassicae* butterflies with impaired olfaction failed to mate, several KO butterflies still mate and deposit fertilized eggs when the oviposition substrate was supplied in our laboratory setting. The mated *Orco* KO butterflies laid a comparable number of fertilized eggs on cabbage plants to WT butterflies which is in line with the results from different moth species (Fandino et al., 2019; Fan et al., 2022; Sun et al., 2023), suggesting oviposition decisions are mostly guided by non-volatile compounds, when no host-plant location is required (van Loon et al., 1992a). Furthermore, the hatching rate of eggs laid by mated *Orco* KO butterflies was similar to that of WT butterflies (Fig S2), indicating that the reduced fecundity in these insects is primarily due to impaired mating behavior and reduced mating frequency, which may result from a lack of pheromonal communication.

When directly placed on their host plant, we found that WT caterpillars gained more weight than *Orco* KO caterpillars (Fig 2A). We speculate that olfaction is needed to evaluate plant tissues and to select different leaves or leaf parts in order to feed efficiently, even when the caterpillar is already on the plant. Secondary metabolites are often unevenly distributed even within a single leaf, and these concentration differences have been shown to influence the performance of different caterpillars (Kester et al., 2002; Yuan et al., 2022). It is conceivable that these localized differences in plant secondary metabolites are also detectable by the herbivore through its olfactory system (Hanson & Dethier, 1972). The difference between WT and KO caterpillars was amplified in the presence of disarmed parasitoids, which were able to attack but not injure the caterpillars, indicating that KO caterpillars were more susceptible to harassment by their natural enemies (Fig 2E). We hypothesize that this further reduction in caterpillar weight was due to the less efficient foraging behavior of the KO caterpillars, which resulted in an extended time period in which the caterpillars were vulnerable to the harassment by the disarmed parasitoids. This harassment would then further reduce the foraging efficiency of the KO caterpillars and thereby exacerbate the effect seen in Fig 2A.

*P. brassicae* caterpillars are most vulnerable to *C. glomerata* during the first two larval instars. Due to their reduced growth rate, KO caterpillars spent a longer time in these vulnerable stages and, therefore, in a natural setting would likely have suffered more strongly from parasitoid attack than WT caterpillars (Brodeur et al., 1996; Haverkamp & Smid, 2020). In addition, caterpillars might have been avoiding plant parts with natural enemies similar to choosing infested plants without parasitoids over infested plants with parasitoids present (Fig 2D). This suggests that caterpillars might be able to use their olfactory system to find enemy-free spaces based on volatiles emitted by the plant or the frass and spit of conspecifics. Similarly, when the caterpillars were exposed to unmanipulated parasitoids, the KO caterpillars were more susceptible and exhibited a higher mortality rate, demonstrating again that olfaction is crucial in larvae to survive under the pressure of natural enemies (Fig 2F). Subsequently, we inferred that the differences in weight gain and survival rate between WT and KO caterpillars are due to different sensitivity to plant volatiles. WT caterpillars can sooner reach a body size in which they are no longer vulnerable to attack by *C. glomerata* by locating an available food source efficiently (Brodeur et al., 1996) . The higher mortality rate of KO caterpillars might, therefore be partly due to a reduced foraging efficiency caused by the lack of olfactory information, which in turn led to an extended period in which the caterpillars were vulnerable to attack by their natural enemies. In addition, the WT caterpillars might have been better able to use olfactory cues to avoid plant parts where conspecifics are under the attack by parasitoids again reducing their mortality in comparison to KO caterpillars (Fig 2D).

Searching for host-plants suitable for feeding and free of natural enemies is largely accomplished via the olfactory system of herbivorous insects (Carrasco et al., 2015; Hu et al., 2020). Without functional ORs, *P. brassicae* caterpillars could still discriminate plant tissues from paper discs; however, *Orco* KO caterpillars preferred the non-host tomato plant over their natural food plant (Fig 2B). We surmise that OR-mediated olfaction is required in caterpillars to locate and identify their host plants, and losing this part of their olfactory receptor repertoire would make it challenging to distinguish host from non-host-plants (Liu et al., 2023). Nonetheless, *Orco* KO caterpillars still retain ORNs expressing ionotropic receptors, which commonly detect acids and amines that have been found to elicit aversion in insect herbivores (Zhang et al., 2019). Fatty acids are present at the edge of both artificially damaged and caterpillar-infested cabbage leaves (Horikoshi et al., 1997), which, in the absence of ORs detecting attractive plant compounds, may have caused avoidance behavior in the KO caterpillar to their natural host-plant. Interestingly, the parasitoid *C. glomerata* also uses the fatty acids produced by the plant to locate *Pieris* caterpillars, and it has been argued that herbivores might avoid these compounds to escape from their natural enemies (Horikoshi et al., 1997; Zhang et al., 2019).

We found that WT caterpillars prefer caterpillar-infested plants free of wasps over caterpillar-infested plants with wasps, which suggests that caterpillars are able to detect certain volatiles that are derived from the interaction with caterpillars and wasps. By analyzing headspace samples of caterpillars, parasitoids as well as the interaction of caterpillars and parasitoids, we successfully identified some chemical compounds which were already known plant volatiles, such as alcohols and aldehydes. However, we did not find any compounds that were specifically abundant in Pb-Cg or Cg. This suggests that a volatile signal serving as a “danger signal” directly emitted by the wasps or caterpillars might not exist, or it might not be detectable with our methods (Fig 3B). Alternatively, indirect cues such as the volatiles emitted from the caterpillar spit during their defensive behavior might have triggered the avoidance behavior of the WT caterpillars to infested plants with natural enemies. When we tested the most abundant compounds in a two-choice behavioral assay against a solvent control, we found that *P. brassicae* WT larvae showed a preference to the plant volatiles (*Z*)-3-hexen-1-ol and (*E*)-2-hexenal (Fig 3D), which are known HIPVs. Previous studies on other species have demonstrated that caterpillars are capable of detecting different HIPVs (Di et al., 2017; de Fouchier et al., 2018). For caterpillars, HIPVs can serve as indicators for the location of potential food plants and the presence of conspecifics (Di et al., 2017; Zhang et al., 2019), but they also attract natural enemies and enhance the risk of being attacked (Bernays, 1997; Ngumbi & Fadamiro, 2012; Yang et al., 2016). Most HIPVs identified in this study were found abundantly in caterpillar saliva that caterpillars spit onto parasitoids to defend against attack. In our study, caterpillars were attracted by HIPVs emitted by plants and by individual HIPVs (Fig 3D). We speculate that caterpillars use low to intermediate concentrations of HIPVs to find suitable host-plants and high concentrations to avoid sites where other caterpillars are under attack by their natural enemies. Therefore, perceiving plant leaf volatiles and approaching the food plants accordingly will increase the herbivores’ feeding efficiency and enhance survival chances. Interestingly, isothiocyanates (ITCs) which are hydrolyzed from glucosinolates after leave damage in crucifer plants, did not elicit strong responses from the butterfly antenna (Fig 1 J and 1K). Because all receptors expressed in the caterpillar are also present in the butterfly and because of the limited number of ORs and ORNs in larvae and low expression levels of ORs (Wang et al., 2024), we found it less likely that ITCs are used by *P. brassicae* caterpillars as foraging cues, even though these compounds are recognized by *Plutella xylostella* ORs (Liu et al., 2020). In our study, volatile cues of wasps were only found in very low quantities and these compounds hardly elicited any behavioral response in the caterpillars; however these chemicals might still play a role in the interaction between the caterpillars and wasps at a close range (Ebrahim et al., 2015).

The way that larval insects react to the potential threat from their natural enemies is still largely unknown for most insects. Our study that exploited CRISPR/Cas9 shows that olfaction is of high significance to caterpillars for locating food sources and likely affects their survival when under the selection pressure from natural enemies. We provided novel insights into the tri-trophic interactions of plants, herbivores and their natural enemies from the perspective of a caterpillar and highlighted the role of olfaction for both foraging and escaping natural enemies.

## Acknowledgements

We appreciate Janneke Bloem for helping with genome editing, and the rearing team at the Laboratory of Entomology for providing insects for our experiments in this study. In addition, we would like to thank the Wageningen Light Microscopy Centre for their support in obtaining the confocal images.

## Author contributions

Conceptualization, Q.W., M.D. and A.H.; Methodology, Q.W., M.J., A.H.; Investigation, Q.W., Y.J., H.S., B.T.W., L.O.G.; Writing - Original Draf, Q.W., A.H.; Writing - Review & Editing, Q.W., L.O.G., M.D., A.H.; Funding Acquisition, M.D., A.H.; Resources: Q.W., M.D., A.H.; Supervision, M.D. and A.H.

## Materials and methods

### Plant and insect rearing

Cabbage plants (*Brassica oleracea* var. gemmifera cv. Cobelius; Brussels sprouts) and tomato plants (*Solanum lycopersicum* cv. moneymaker) were planted in individual pots and grown for four weeks after germination before being used in the experiments or to feed the caterpillars. The cabbage plants were reared under glasshouse conditions at 22 ± 3 °C, RH 50-80 % and 16 h light: 8 h dark cycles. *Pieris brassicae* caterpillars and butterflies as well as *C. glomerata* wasps were collected from laboratory colonies maintained at the Laboratory of Entomology, Wageningen University & Research, the Netherlands. The caterpillars were reared on Brussels sprouts plants at 22 ± 3 °C, RH 25-35 % with 14 h light: 10 h dark cycles. The adults were supplied with 10 % sugar water as nutrient under the same environmental conditions. After successful gene editing, KO insects were transferred and reared in a glass house compartment under conditions at 25 ± 3 °C, RH 25-35 %. Several KO caterpillars and butterflies from each generation were randomly collected for genotype screening to ensure that homozygosity remains. *Cotesia glomerata* adults were fed with organic honey, *P. brassicae* second-instar larvae (L2) were supplied to the parasitoid wasps for parasitization to maintain the colony. The wasp cocoons were collected in Petri dishes and incubated at 22 °C, RH 50-70 % with the same photoperiod as that of caterpillars. All the subsequent comparisons between WT and KO in this study were conducted in the glass house compartment under the rearing condition of KO insects unless specified.

### *Orco* knockout

The second exon of *Orco* was targeted to knockout the gene. The sgRNA was designed with Geneious Prime 2021 (Geneious, New Zealand) by searching N20 target + NGG PAM sequences in the second exon, sgRNA primers were selected according to activity scorings (Doench et al., 2016). The designated sgRNA off-target regions were evaluated by searching against the genome by est2genome function with a loose 50-score cut-off by using Exonerate 2.0 (Slater & Birney, 2005) and CHOPCHOP v3 (Labun et al., 2019). The sgRNAs that had no off-target sites in coding areas were selected. sgRNA templates were prepared by mixtures of 50 µL Q5 high-fidelity 2 × master mix (NEB, USA), 5 µL of the following forward primers: sgRNA1: ATTTAGGTGACACTATACATGTCAACTCTACTTAGGAGTTTTAGAGCTAGAAATA GCAAG; sgRNA2: ATTTAGGTGACACTATACGATGAAGTAAACGAACTCAGTTTTAGAGCTAGAAATAGCAAG; 5 µL constant reverse primer: AAAAGCACCGACTCGGTGCCACTTTTTCAAGTTGATAACGGACTAGCCTTATTTTAACTTGCTATTTCTAGCTCTAAAAC and 40 µL H_2_O. The mixture was incubated at 98 °C for 2 min; followed by 38 cycles of 98 °C for 20 s, 65 °C for 10 s and 72 °C for 10 s; finalized at 72 °C for 5 min. The PCR products were checked by 1.0 % gel electrophoresis and purified by QIAGEN PCR purification kit (QIAGEN, Germany) according to the manufacturer’s instructions. sgRNA was synthesized by a reaction system of 2 µL 10 × SP6 enzyme mix (Invitrogen, USA), 2 µL 10 × SP6 reaction buffer, 2 µL each of ATP, UTP, GTP and CTP, 200 ug sgRNA template and add RNase-free H_2_O up to 20 µL. The mixture was incubated at 37 °C for 4 hours, followed by adding 1 µL Turbo DNase and incubating at 37 °C for 15 min. The reaction products were purified by Monarch RNA cleanup kit (NEB, USA). The concentration of sgRNA was determined by DeNovix (DeNovix, USA). sgRNA was mixed with Cas9 protein (NEB, USA) and incubated at 25 °C for 10 min. Newly laid eggs, not older than half an hour, were collected from our laboratory colony and were fixed on glass slides. The sgRNA/Cas9 mixture was colored with 1 µL food dye to stain the eggs as an indicator that injection with a Femto-jet (Eppendorf, Germany) was successful. The injected eggs were incubated at 25 °C for 4 to 5 days until hatching. The caterpillars were reared until the emergence of the butterflies, a leg of each butterfly was cut for genotyping. gDNA of the leg samples was extracted with MyTaq Extract-PCR Kit (Bioline, Germany). Mutant screening was performed by PCR with 15 µL 2 × MyTaq HS Red mix, 2 µL forward primer (TCTGGCTTCGGTATTACATTTC), 2 µL reverse primer (CTTTTATGGCGTGTTTTATTTG), 2 µL gDNA template and 9 µL H_2_O. The PCR products were checked by gel electrophoresis and sequenced with the same primers by Eurofins (Eurofins, the Netherlands).

### ORN staining and axon tracing

L3 caterpillars (n = 6 for WT and n = 5 for KO) were anesthetized on ice and decapitated. Heads were cut along the midline with micro-scissors to facilitate penetration of the fixative and antibodies. The collected heads were immersed in freshly prepared 4 % formaldehyde in 0.1 M phosphate buffer at pH 7.3 and fixed overnight at 4 °C. The fixed samples were rinsed in 70 % ethanol, followed by incubation in a mixture of 96 % ethanol: 30 % hydrogen peroxide at 1:1 ratio for 7 days at 4 °C. The incubated samples were then dehydrated in a graded series of ethanol, degreased in xylene for two mins, rehydrated and incubated in 0.1 M phosphate buffered saline (Oxoid, Dulbecco A) with 0.5% triton X-100 (PBS-T) followed by four washes in PBS-T for 15 min each. Preincubation of head samples was performed in 10 % normal goat serum (Bio-connect services, The Netherlands) in PBS-T (PBS-T-NGS) for 1 hour, and then incubated in rabbit anti-Orco antibody which specificity was tested previously (Nolte et al., 2016) diluted in 1: 500 in PBS-T-NGS for three days at 4 °C, followed by 6 washes in PBS-T for 30 min each. Samples were subsequently incubated in goat anti-rabbit conjugated to Alexa fluor 488 (Thermo Fisher Scientific, USA) diluted 1:100 and TO-PRO-3 iodide (Thermo Fisher Scientific, USA) diluted 1:1000 in PBS-T-NGS for two days at 4 °C. Heads were washed two times in PBS-T for 30 mins and once overnight at 4 °C, following with two 1-hour PBS washes at room temperature. The samples were then dehydrated with a graded series of ethanol, cleared in xylene and mounted in DPX. Confocal microscopy was performed with a Leica Stellaris 5-DIM 8 confocal microscope (Leica, Germany), using a 63× oil-immersion plan APO objective NA1.4. and a spectrally flexible white light laser for excitation of the two fluorophores using the pre-sets for Alexa fluor 488 and To-Pro-3 iodide.

L4 caterpillars of similar head size were anesthetized on ice and immobilized in clay. The tip of each antenna was peeled, and a layer of petroleum jelly (Vaseline) was applied around the antenna. A 0.5 µL drop of 2.5 % biotin-dextran solution was added to the cut and covered with petroleum jelly, then incubated at 4 °C overnight. Brains were dissected in PBS, fixed in 4 % formaldehyde in PBS at 4 °C overnight, dehydrated in xylene and rehydrated in PBS-T, followed by three washes in PBS-T for 2 hours each. Brains were then incubated in 10 % NGS in PBS-T for 1 hour, followed by incubation in 1:200 streptavidin 488 and 1:1000 To-Pro iodide in PBS-T at 4 °C for three days. After further rinsing in PBS-T overnight, brains were mounted in DPX. Whole-mount samples were scanned using confocal microscopy with settings similar to those used for ORN staining.

### Electroantennography (EAG)

A panel of 54 chemicals (Table S1) including esters, alcohols, aldehydes, alkenes, isothiocyanates (ITCs), heterocyclics and nitriles were selected from the literature (van Loon et al., 1992b; Liu et al., 2020; Zhu et al., 2015; Bourne et al., 2023) and our current volatile identification data, to investigate the EAG responses of both KO and WT butterfly antennae (n =15-18) of both sexes. Filter papers loaded with 10 µL of selected chemicals, diluted in paraffin oil (10^-2^ v/v), were inserted into Pasteur pipettes, clean filter paper loaded with 10 µL paraffin oil was employed as negative control. The left antennae of butterflies were excised at the basal end and the distal tip was also removed for better conductivity. Antennae were placed into glass capillaries filled with EAG Ringer solution. Glass capillaries were placed on Ag-AgCl wires connected to a ground electrode and a 10× high-impedance DC amplifier (Ockenfels Syntech, Germany). The electrical signals were transformed using an IDAC-232 (Ockenfels Syntech, Germany) analog-digital converter connected to a personal computer. Signals were finally recorded using the software EAG Pro (Ockenfels Syntech, Germany). The first puff of each chemical was abandoned before the first round of testing. Antennae were exposed to the chemicals in random order and were given intervals of at least 20 s for recovering until the baseline stabilized.

### Female oviposition behavior

A mating pair of newly emerged butterflies (n = 11 for WT and n = 14 for KO) were put in a cage and supplied with a four-week-old Brussels sprouts cabbage plant as oviposition substrate and sugar water as nutrients. The numbers of eggs were counted on a daily basis for seven days. The female butterflies were then collected and dissected to compare mating frequency between WT and KO females. Mating frequency was determined by the number of spermatophores dissected from the females’ abdomens. The eggs remained on the plants for another four to five days to allow hatching. The number of caterpillars hatched enabled us to compare the number of fertilized eggs and hatching rate between the two butterfly genotypes.

### Caterpillar performance on plants

Caterpillar performance on cabbage plants was evaluated by three different treatments:

1. For each genotype, a group of ten L1 caterpillars (n = 17) was placed on one leaf of a four-week-old cabbage plant. Caterpillars were reared for ten days, and more plants were provided when needed. The caterpillars were then collected for weighing to compare the growth between the two genotypes.
2. The growth of both genotypes of caterpillars (n = 8) was further compared by rearing in the presence of *C. glomerata* parasitoid wasps from which the ovipositor had been removed (hereafter: “disarmed”). Groups of ten L1 caterpillars were placed on individual four-week-old cabbage plants as described above. The parasitoid wasps were anesthetized on a CO_2_ plate and the ovipositor was removed with fine tweezers to ensure that the parasitoid wasps can interact with the caterpillars but cannot parasitize them. The activity of disarmed parasitoids was observed for several minutes to ensure that they would behave similarly to unmanipulated parasitoids. Two disarmed parasitoid wasps were released in each rearing cage 24 hours after the caterpillars had been placed on the plants. The disarmed parasitoid wasps were replaced by newly disarmed parasitoids every day, the caterpillars were reared on the plants for ten days and then collected for weighing.
3. To compare the survival rate of caterpillars (n = 7 groups of ten caterpillars) on the plants under the threat of *C. glomerata* parasitoid wasps, a group of ten L1 caterpillars was placed on a leaf of a four-week-old cabbage plant two days prior to placing two female parasitoid wasps. The parasitoid wasps were replaced every day to ensure that caterpillars were always exposed to natural enemies. The caterpillars were collected after ten days to count the number that survived; the collected caterpillars were further reared until they pupated or *C. glomerata* larvae emerged.

### Larval host-plant seeking behavior

Caterpillar (n = 54-73) food-plant seeking behavior was evaluated in a greenhouse under the same conditions in which the caterpillars were reared. Plant leaf discs or green paper discs (Trophée 1224 “Forest Green”, Clairefontaine, France), with 12 mm diameter were placed at opposite positions along the edge of a 188 mm-diameter Petri dish. A single L3 caterpillar was then placed at the center of the arena. Each caterpillar was observed for 10 min, the first choice that the caterpillars made and the time spent were recorded. The caterpillars were provided with a choice between either (i) a cabbage leaf disc and a paper disc, (ii) a tomato leaf disc and a paper disc, or (iii) a cabbage leaf disc and a tomato leaf disc. A choice was recorded when caterpillars began feeding on the cabbage leaf disc or contacted the tomato leaf / green paper discs. New discs were used for every caterpillar and the positions of discs were exchanged after every ten tests.

To further test the host-plant locating behavior of L3 caterpillars, we investigated the behaviors of caterpillars in a Y-tube olfactometer (diameter 35 mm) with a built-in black Y-shaped wire, allowing caterpillars to crawl along the wire and make a choice smoothly in the laboratory. Each arm was provided with a 1.1 L/min inlet airflow which had been purified by charcoal and humidified by water before entering the treatment jars. Healthy plants were untreated, infested plants were infested with 30 early L3 caterpillars for 24 hours and with caterpillars on plants while testing. Differently treated plants were placed in jars to test caterpillar behaviors. The caterpillars had been starved for approximately 5 hours before testing. Each caterpillar was tested and observed for 5 min, caterpillar behavior was monitored by a camera. The position of inlet airflow was changed every five tests to avoid position bias.

### Volatile analysis

Volatiles were collected from L3 *P. brassicae* caterpillars, three-day-old female *C. glomerata* wasps, *P. brassicae* caterpillars and *C. glomerata* wasps together, caterpillar frass collected from L3 caterpillars and caterpillar saliva also collected from L3 caterpillars using glass capillaries following Mattiacci et al. (Mattiacci et al., 1994). Fresh frass and saliva samples were stored in a freezer at -20 °C and thawed overnight at room temperature before volatile collection. Headspace collection of volatiles was performed from five treatments: 20 female wasps (Cg) in a 10 mL glass vial (n =12); 20 L3 caterpillars (Pb) in a 10 mL glass vial (n =10); 20 L3 caterpillars and 2 female wasps (Pb-Cg) in a 10 mL glass vial (n =14); 100 mg caterpillar frass (Pb-Fr) in a 1.5 mL glass vial (n =10); and 50 µL caterpillar saliva (Pb-S) in a 1.5 mL glass vial (n =12). Clean empty glass vials (10 mL and 1.5 mL) were used as a negative control. Volatiles were collected through dynamic headspace sampling using Tenax TA adsorbent material (20/35 mesh; Camsco, USA). Synthetic air (Air Synthetic 4.0 Monitoring from Linde Gas, the Netherlands) at 110 mL min^-1^ to the samples in 10 mL vials and 55 mL min^-1^ to the samples in 1.5 mL vials were constantly supplied as a carrier for the volatiles, while simultaneously the volatiles were trapped by drawing air at 100 and 50 mL min^-1^, respectively, through a stainless-steel tube filled with 200 mg Tenax TA for 2 h.

The collected volatiles were thermally released from the Tenax TA adsorbent using an Ultra 50:50 thermal desorption unit (Markes, UK) at 250 °C for 10 min under a helium flow of 20 mL min^-1^, while simultaneously re-collecting the volatiles in a thermally cooled universal solvent trap: Unity (Markes, UK) at 0 °C. Once the desorption process was completed, the volatile compounds were released from the cold trap by ballistic heating at 40°C s^-1^ to 280 °C, which was then kept for 10 min, while all the volatiles were transferred to a ZB-5MS analytical column (30 mL × 0.25 mm ID × 1 mm F.T. with 10 m built-in guard column (Phenomenex, USA), placed inside the oven of a Thermo Trace GC Ultra (Thermo Fisher Scientific, USA), for separation of volatiles. The GC oven temperature was initially held at 40 °C for 2 min and was immediately raised at 6 °C min^-1^ to a final temperature of 280°C, where it was kept for 4 min under a constant helium flow of 1 mL min^-1^. For the detection of volatiles, a Thermo Trace DSQ quadrupole mass spectrometer (Thermo Fisher Scientific, USA) coupled to the GC was operated in an electron impact ionization (EI) mode at 70 eV in a fullscan mode with a mass range of 35 - 400 amu at 4.70 scans s^-1^. The MS transfer line and ion source were set at 275 and 250 °C, respectively.

Automated baseline correction, peak selection (S/N > 3) and alignments of all extracted mass signals of the raw data were processed following an untargeted metabolomic workflow using MetAlign software, producing detailed information on the relative abundance of mass signals representing the available metabolites (Lommen, 2009). This was followed by the reconstruction of the extracted mass features into potential compounds using the MSClust software through data reduction by means of unsupervised clustering and extraction of putative metabolite mass spectra (Tikunov et al., 2012). Tentative identification of volatile metabolites was based on a comparison of the reconstructed mass spectra with those in the NIST 2008 and Wageningen Mass Spectral Database of Natural Products MS libraries, as well as experimentally obtained linear retention indices (LRIs).

### Caterpillar response to individual chemical compounds

L3 caterpillars (n = 31-50 for each compound, each genotype) were tested in a custom-made set-up consisting of 25 separate arenas in the laboratory (Fig 3C). Each arena allowed caterpillars to move freely along two channels of 90 mm located in opposite directions of a central opening, which permitted the caterpillar to crawl into the arena. At the end of the arenas, airflows are introduced to deliver volatile chemicals. The airflow was sucked out through the channel below the central opening simultaneously. Single caterpillars were placed in the central well of the individual arenas before the test. Caterpillars were given a choice between a selected volatile chemical of interest (Table S2) and paraffin oil. Input odors were delivered at 1.1 L min^-1^ for both chemicals and solvent controls, the odors were sucked out at 2.2 L min^-1^ at the same time from the center. The arena was cleaned by ethanol and dried by high-pressure air. The position of chemicals was exchanged after each test. The behavior of caterpillars was recorded for 10 min at 100 frames per minute using a Canon TV lens JF16mm (Canon, Japan). Behavioral data was extracted by EthoVision XT 11.5 software (Noldus, The Netherlands). Data were removed from the dataset when the caterpillar stayed in the arena for less than 10 seconds. All tests were performed at room temperature.

### Statistical analysis

Unless stated otherwise, statistical tests were performed in R (version 4.4.0; R Core Team, 2016) in combination with RStudio (Posit, USA). Data sets were tested for normal distribution using a Shapiro– Wilk test and for equal variances using Levene’s test. The difference of glomerular number between WT and *Orco* KO caterpillars was tested with a Wilcoxon rank-sum test. Electroantennographic responses between the two genotypes were tested statistically by using the Student’s t-test when the values were normally distributed and had homogenous variance or tested by using the Kruskal-Wallis rank-sum test when the requirements for the t-test were not met. The difference in butterfly mating frequency was analyzed by using the Wilcoxon rank-sum test; differences in oviposition dynamics were tested with a generalized linear model (GLM) with a negative binominal distribution. The development differences between WT and KO caterpillars on full plants and under the thread of disarmed parasitoids were tested using a Student’s t-test, and survival rate difference between the two genotypes was tested using a GLM with a beta-binomial distribution. Caterpillar host-plant selection behaviors in Petri dishes and in the Y-tube were analyzed with a Chi-square test. Differences of the relative time spent by the caterpillars between the control side and the odor side of the multichannel arenas were compared by a non-parametric Wilcoxon rank-sum test. Principal component analysis (PCA) of volatile blends was achieved by MetaboAnalyst (Pang et al., 2024). Data were normalized by logarithmic transformation and scaled in auto mode. Ward’s method and Euclidean distance were employed in the hierarchical clustering heatmap by default settings without clustering samples. In addition, the abundance of volatiles as peak heights was imported into SIMCA-P 17 statistical software (Umetrics, Sweden), the data was analyzed by multivariate data analysis (MVDA). Supervised orthogonal partial least-squares discriminant analysis (OPLS-DA) was employed to compare and correlate treatment groups.

## Supporting information

**Fig S1. *Pieris brassicae* caterpillar development in a Petri dish environment.** X axis indicates the date since caterpillars have been deposited in the Petri dish. Y axis indicates the weight of caterpillars (Ln Weight). Caterpillar weight (n = 10 for both genotypes) in this figure is presented by the mean of a group of caterpillars weight in each Petri dish on day 6.

**Fig S2. Egg-hatching rates by *wildtype* (n=11), *Orco^-/-^*_fert (n=5) and *Orco^-/-^*_unfert (n=9) butterflies.** *Orco^-/-^*_fert represents mated Orco female butterflies, *Orco^-/-^*_unfert represents unmated female butterflies. Differences were tested by using Student’s t-test.

**Fig S3. Mating frequency of *Pieris brassicae* butterflies.** The boxplots indicate the number of spermatophores in WT female butterflies (n = 11) and in KO butterflies (Orco-/-) (n = 14), the difference was tested by using Wilcoxon rank-sum test. WT and Orco KO butterfly spermatophores are indicated in orange and blue respectively.

**Fig S4. *Pieris brassicae* wildtype and *Orco* knockout (*Orco ^-/-^*) butterfly oviposition dynamics.** The orange line indicates the number of eggs per female (mean ± SE) laid by wildtype (WT) butterflies (n = 11), the blue line indicates the number of eggs laid by Orco knockout (KO) butterflies (n = 14). Differences were analyzed using a GLM with negative binominal distribution, difference test results are shown in the line chart.

**Fig S5. Overview of volatile blends of *Pieris brassicae* caterpillars, *Cotesia glomerata* parasitoid wasps and caterpillars-parasitoid in interaction.** (A), OPLS-DA (Orthogonal Projection to Latent Structures Discriminant Analysis) two-dimensional score plot of treatment groups: Cg, *Cotesia glomerata* female parasitoid wasps (n = 12), based on their volatile content; Pb, *Pieris brassicae* caterpillars (n = 10); Pb-Cg, *P. brassicae* caterpillars in the presence of *C. glomerata* female parasitoid wasps (n = 14. (B), Loading plot showing the contribution of each identified volatile compound to the separation of the different treatments. Volatiles closer to the treatment in the plot indicate higher correlation. Numbers in the loading plot refer to the volatile compounds listed in Supporting information Table S3.

**Fig S6. Overview of volatile blends of *Pieris brassicae* caterpillars and caterpillars-parasitoid (*Cotesia glomerata*) in interaction.** (A), OPLS-DA (Orthogonal Projection to Latent Structures Discriminant Analysis) two-dimensional score plot of treatment groups: Pb, *Pieris brassicae* caterpillars (n = 10); Pb-Cg, *P. brassicae* caterpillars in the presence of *C. glomerata* female parasitoid wasps (n = 14), based on their volatile content. (B), Loading plot showing the contribution of each identified volatile compound to the separation of the different treatments. Volatiles closer to the treatment in the plot indicate higher correlation. Numbers in the loading plot refer to the volatile compounds listed in Supporting information Table S3.

**Fig S7. Overview of volatile blends of *Pieris brassicae* caterpillar spit and caterpillars-parasitoid wasp (*Cotesia glomerata*) in interaction.** (A), OPLS-DA (Orthogonal Projection to Latent Structures Discriminant Analysis) two-dimensional score plot of treatment groups: P. brassicae spit (n = 12), Pb-Cg, *P. brassicae* caterpillars in the presence of *C. glomerata* female parasitoid wasps (n = 14), based on their volatile content. (B), Loading plot showing the contribution of each identified volatile compound to the separation of the different treatments. Volatiles closer to the treatment in the plot indicate higher correlation. Numbers in the loading plot refer to the volatile compounds listed in Supporting information Table S3.

**Fig S8. Overview of volatile blends of *Pieris brassicae* caterpillar spit and *Pieris brassicae* caterpillar frass.** (A), OPLS-DA (Orthogonal Projection to Latent Structures Discriminant Analysis) two-dimensional score plot of treatment groups: Pb-Fr, *P. brassicae* caterpillar frass (n = 10); and Pb-S, *P. brassicae* spit (n = 12), based on their volatile content. (B), Loading plot showing the contribution of each identified volatile compound to the separation of the different treatments. Volatiles closer to the treatment in the plot indicate higher correlation. Numbers in the loading plot refer to the volatile compounds listed in Supporting information Table S3.

**Fig S9. Overview of volatile blends of *Pieris brassicae* caterpillars, *Pieris brassicae* caterpillar spit and *Pieris brassicae* caterpillar frass.** (A), OPLS-DA (Orthogonal Projection to Latent Structures Discriminant Analysis) two-dimensional score plot of treatment groups: Pb, *Pieris brassicae* caterpillars (n = 10); Pb-Fr, *P. brassicae* caterpillar frass (n = 10); and Pb-S, *P. brassicae* spit (n = 12), based on their volatile content. (B), Loading plot showing the contribution of each identified volatile compound to the separation of the different treatments. Volatiles closer to the treatment in the plot indicate higher correlation. Numbers in the loading plot refer to the volatile compounds listed in Supporting information Table S3.

**Fig S10. Heatmaps of WT caterpillar movement in response to the tested chemicals.** The chemical compounds are indicated in the figures and color legend indicates the time (s) that each caterpillar stayed at certain locations.

**Fig S11. Heatmaps of Orco KO caterpillar movement in response to the tested chemicals.** The chemical compounds are indicated in the figures and color legend indicates the time (s) that each caterpillar stayed at certain locations.

**Table S1. Chemical compounds that were used for the electroantennographical test.**

**Table S2. Chemical compounds that were used for the behavioral test in multi-channel arena.**

**Table S3. Volatile compounds detected in the headspace of the different treatment samples.** Cg, *Cotesia glomerata* female parasitoid wasps (n = 12); Pb, *Pieris brassicae* caterpillars (n = 10); Pb-Cg, *P. brassicae* caterpillars in the presence of *C. glomerata* female parasitoid wasps (n = 14); Pb-Fr, *P. brassicae* caterpillar frass (n = 10); and Pb-S, *P. brassicae* spit (n = 12), are listed. Relative amounts of volatiles are presented as average peak height (SE)/10^4^. The volatiles are listed according to their elution order in a chromatographic window. Volatiles with Variable Importance in the Projection (VIP) scores equal to or higher than 1.0 (presented in bold), are considered important in separating the different treatment groups of the given analysis. Significant differences among the five treatments detected by Kruskal-Wallis test with Dunn’s post-hoc test for multiple comparisons are indicated by letters in the table. Different letters indicate significant difference (P < 0.05). NF, not found.

**Table S4. Volatiles used in separating *Cotesia glomerata* female wasps (Cg, n = 12), *Pieris brassicae* caterpillars (Pb, n = 10) and the interaction of *P. brassicae* caterpillars with *C. glomerata* female wasps (Pb-Cg, n = 14) sample treatments (Fig S5).** Volatiles are listed according to ranking order of their Variable Importance in the Projection (VIP) scores values, where those with VIP scores of equal to or higher than 1.0, are considered important in separating the treatment groups of the given analysis.

**Table S5. Volatiles used in separating *Pieris brassicae* caterpillars (Pb, n = 10) and the interaction of *P. brassicae* caterpillars with *Cotesia glomerata* female wasps (Pb-Cg, n = 14) sample treatments (Fig S6).** Volatiles are listed according to ranking order of their Variable Importance in the Projection (VIP) scores values, where those with VIP scores of equal to or higher than 1.0, are considered important in separating the treatment groups of the given analysis.

**Table S6. Volatiles used in separating the interaction of *Pieris brassicae* caterpillars with *Cotesia glomerata* female wasps (Pb-Cg, n = 14) and *P. brassicae* caterpillar spit (Pb-S, n = 12), sample treatments (Fig S7).** Volatiles are listed according to ranking order of their Variable Importance in the Projection (VIP) scores values, where those with VIP scores of equal to or higher than 1.0, are considered important in separating the treatment groups of the given analysis.

**Table S7. Volatiles used in separating the *Pieris brassicae* caterpillar spit (Pb-S, n = 12), and *P. brassicae* caterpillar frass (Pb-Fr, n = 10) sample treatments (Fig S8).** Volatiles are listed according to ranking order of their Variable Importance in the Projection (VIP) scores values, where those with VIP scores of equal to or higher than 1.0, are considered important in separating the treatment groups of the given analysis.

**Table S8. Volatiles used in separating the *Pieris brassicae* caterpillars (Pb, n = 10), the interaction of *P. brassicae* caterpillars with *Cotesia glomerata* female wasps (Pb-Cg, n = 14), and *P. brassicae* caterpillar frass (Pb-Fr, n = 10) sample treatments (Fig S9).** Volatiles are listed according to ranking order of their Variable Importance in the Projection (VIP) scores values, where those with VIP scores of equal to or higher than 1.0, are considered important in separating the treatment groups of the given analysis.

