## Supporting information for "Loss of olfaction reduces caterpillar performance and increases susceptibility to a natural enemy"

https://doi.org/10.7554/eLife.105585.1

Qi Wang [1], Yufei Jia [1], Hans M. Smid [1], Berhane T. Weldegergis [1], Liana O. Greenberg [2], Maarten Jongsma [3], Marcel Dicke [1], Alexander Haverkamp [1,4]*

[1] Laboratory of Entomology, Wageningen University & Research, Wageningen, the Netherlands

[2] Biosystematics Group, Wageningen University & Research, Wageningen, the Netherlands

[3] Business Unit Bioscience, Wageningen University & Research, Wageningen, the Netherlands

[4] Lead contact

**Table of Contents**

Table S1. Chemical compounds that were used for the electroantennographical test.

Table S2. Chemical compounds that were used for the behavioral test in multi-channel arena.

Table S3. Volatile compounds detected in the headspace of the different treatment samples.

Table S4. Volatiles used in separating *Cotesia glomerata* female wasps (Cg, n = 12), *Pieris brassicae* caterpillars (Pb, n = 10) and the interaction of *P. brassicae* caterpillars with *C. glomerata* female wasps (Pb-Cg, n = 14) sample treatments (Fig S5).

Table S5. Volatiles used in separating *Pieris brassicae* caterpillars (Pb, n = 10) and the interaction of *P. brassicae* caterpillars with *C. glomerata* female wasps (Pb-Cg, n = 14) sample treatments

Table S6. Volatiles used in separating the interaction of *Pieris brassicae* caterpillars with *Cotesia glomerata* female wasps (Pb-Cg, n = 14) and *P. brassicae* caterpillar spit (Pb-S, n = 12), sample treatments (Fig S7).

Table S7. Volatiles used in separating the *Pieris brassicae* caterpillar spit (Pb-S, n = 12), and *P. brassicae* caterpillar frass (Pb-Fr, n = 10) sample treatments (Fig S8).

Table S8. Volatiles used in separating the *Pieris brassicae* caterpillars (Pb, n = 10), the interaction of *P. brassicae* caterpillars with *Cotesia glomerata* female wasps (Pb-Cg, n = 14), and *P. brassicae* caterpillar frass (Pb-Fr, n = 10) sample treatments (Fig S9).

Fig S1. *Pieris brassicae* caterpillar development in a Petri dish environment.

Fig S2. Egg-hatching rates by *wildtype* (n=11), *Orco*$^{-/-}$_fert (n=5) and *Orco*$^{-/-}$_unfert (n=9)

Fig S3. Mating frequency of Pieris brassicae butterflies.

Fig S4. *Pieris brassicae wildtype* and *Orco* knockout (*Orco*$^{-/-}$) butterfly oviposition dynamics.

Fig S5. Overview of volatile blends of *Pieris brassicae* caterpillars, *Cotesia glomerata* parasitoid wasps and caterpillars-parasitoid in interaction.

Fig S6. Overview of volatile blends of *Pieris brassicae* caterpillars and caterpillars-parasitoid (*Cotesia glomerata*) in interaction.

Fig S7. Overview of volatile blends of *Pieris brassicae* caterpillar spit and caterpillars-parasitoid wasp (*Cotesia glomerata*) in interaction.

Fig S8. Overview of volatile blends of *Pieris brassicae* caterpillar spit and *P. brassicae* caterpillar

Fig S9. Overview of volatile blends of *Pieris brassicae* caterpillars, *P. brassicae* caterpillar spit and *P. brassicae* caterpillar frass.

Fig S10. Heatmaps of WT caterpillar movement in response to the tested chemicals.

Fig S11. Heatmaps of *Orco* KO caterpillar movement in response to the tested chemicals.

**Table S1. Chemical compounds that were used for the electroantennographical test**

| Chemical compound | CAS number | Purity | Manufacturer |
|---|---|---|---|
| Acetic acid | 64-19-7 | ⩾ 99.0% | Sigma-Aldrich |
| Benzoic acid | 65-85-0 | ⩾99.5% | Sigma-Aldrich |
| Isovaleric acid | 503-74-2 | 99.00% | Sigma-Aldrich |
| n-Caproic acid | 142-62-1 | ⩾ 99.0% | Sigma-Aldrich |
| (*E*)-Anethole | 4180-23-8 | 99.00% | Sigma-Aldrich |
| 1,8-Cineole | 470-82-6 | 99.00% | Sigma-Aldrich |
| 1-Hexanol | 111-27-3 | 98.00% | Fluka |
| 1-Methoxy-2-propanol | 107-98-2 | ⩾ 99.5% | Sigma-Aldrich |
| 1-Octen-3-ol | 3391-86-4 | 98.00% | Sigma-Aldrich |
| 1-Pentanol | 71-41-0 | ⩾ 99.0% | Sigma-Aldrich |
| 1-Penten-3-ol | 616-25-1 | 99.00% | Sigma-Aldrich |
| 3-Methyl-1-butanol | 123-51-3 | ⩾ 99.0% | Sigma-Aldrich |
| 3-Octanol | 589-98-0 | 99.00% | Sigma-Aldrich |
| 3-Pentanol | 584-02-1 | 98.00% | Sigma-Aldrich |
| (*Z*)-2-Penten-1-ol | 1576-95-0 | 95.00% | Sigma-Aldrich |
| (*Z*)-3-Hexen-1-ol | 928-96-1 | 98.00% | Sigma-Aldrich |
| Geraniol | 106-24-1 | 98.00% | Sigma-Aldrich |
| Linalool | 78-70-6 | 97.00% | Sigma-Aldrich |
| Phenylethyl alcohol | 8-12-1960 | ⩾99.0 % | Fluka |
| (*E*)-2-Hexen-1-ol | 928-95-0 | ⩾ 95.0% | Sigma-Aldrich |
| 1-Nonanal | 124-19-6 | 95.00% | Sigma-Aldrich |
| 2-Methylbutanal | 96-17-3 | 95.00% | Sigma-Aldrich |
| 3-Methylbutanal | 590-86-3 | ⩾ 98.5% | Fluka |
| Benzaldehyde | 100-52-7 | ⩾99.0% | Fluka |
| (*Z*)-2-Pentenal | 1576-87-0 | 95.00% | Sigma-Aldrich |
| Heptaldehyde | 111-71-7 | 95.00% | Sigma-Aldrich |
| Hexanal | 66-25-1 | 98.00% | Sigma-Aldrich |
| Phenylacetaldehyde | 122-78-1 | ⩾95.0% | Fluka |
| (*E*)-2-Hexenal | 6728-26-3 | 98.00% | Sigma-Aldrich |
| Nonane | 111-84-2 | 99.00% | Sigma-Aldrich |
| α-Pinene | 80-56-8 | 98.00% | Sigma-Aldrich |
| β-Caryophyllene | 87-44-5 | ⩾ 98.0% | Sigma-Aldrich |
| β-Pinene | 127-91-3 | ⩾ 95.0% | Sigma-Aldrich |
| Limonene | 5989-27-5 | 97.00% | Sigma-Aldrich |
| Myrcene | 123-35-3 | ⩾ 99.0 | Sigma-Aldrich |
| (*Z*)-3-Hexenyl acetate | 3681-71-8 | 98.00% | Sigma-Aldrich |
| Hexyl acetate | 142-92-7 | 99.00% | Sigma-Aldrich |
| Methyl salicylate | 119-36-8 | ⩾ 98.0% | Sigma-Aldrich |
| Pentyl acetate | 628-63-7 | 99.00% | Sigma-Aldrich |
| Benzothiazole | 95-16-9 | ⩾ 96.0% | Sigma-Aldrich |
| Indole | 120-72-9 | ⩾ 99.0% | Sigma-Aldrich |
| Allyl ITC | 7-6-1957 | ⩾ 95.0% | Sigma-Aldrich |
| Benzyl ITC | 622-78-6 | 98.00% | Sigma-Aldrich |
| Butyl ITC | 592-82-5 | 99.00% | Sigma-Aldrich |
| Methyl ITC | 556-61-6 | 97.00% | Fluka |
| Phenyl ITC | 103-72-0 | 98.00% | Sigma-Aldrich |
| 2,3-Butanedione | 431-03-8 | 97.00% | Fluka |
| 3-Hydroxy-2-butanone | 513-86-0 | ⩾ 98.0% | Sigma-Aldrich |
| 3-Octanone | 106-68-3 | ⩾ 98.0% | Sigma-Aldrich |
| 3-Pentanone | 96-22-0 | ⩾ 99.0% | Sigma-Aldrich |
| 3-Butenenitrile | 109-75-1 | 98.00% | Sigma-Aldrich |
| Benzyl cyanide | 140-29-4 | 98.00% | Sigma-Aldrich |
| Dimethyl disulfide | 624-92-0 | ⩾ 99.0% | Sigma-Aldrich |
| Paraffin oil | 8012-95-1 | - | Sigma-Aldrich |

**Table S2. Chemical compounds that were used for the behavioral test in multi-channel arena.**

| Chemical compound | CAS number | Purity | Manufacturer |
|---|---|---|---|
| 1-Hexanol | 111-27-3 | 98.00% | Fluka |
| 1-Methoxy-2-propanol | 107-98-2 | ≥ 99.5% | Sigma-Aldrich |
| 1-Penten-3-ol | 616-25-1 | 99.00% | Sigma-Aldrich |
| 2,3-Butanedione | 431-03-8 | 97.00% | Fluka |
| 3-Methylbutanal | 123-51-3 | ≥ 98.5% | Fluka |
| Benzyl cyanide | 140-29-4 | 98.00% | Sigma-Aldrich |
| Dimethyl disulfide | 624-92-0 | ≥ 99.0% | Sigma-Aldrich |
| Pentanal | 110-62-3 | ≥ 99.0% | Sigma-Aldrich |
| (*E*)-2-Hexen-1-ol | 928-95-0 | ≥ 95.0% | Sigma-Aldrich |
| (*Z*)-2-Penten-1-ol | 1576-95-0 | 95.00% | Sigma-Aldrich |
| (*Z*)-3-Hexen-1-ol | 928-96-1 | 98.00% | Sigma-Aldrich |
| Linalool | 78-70-6 | 97.00% | Sigma-Aldrich |
| (*Z*)-3-Hexenyl acetate | 3681-71-8 | 98.00% | Sigma-Aldrich |
| Hexanal | 66-25-1 | 98.00% | Sigma-Aldrich |
| (*E*)-2-Hexenal | 6728-26-3 | 98.00% | Sigma-Aldrich |

**Table S3. Volatile compounds detected in the headspace of the different treatment samples.** Cg, *Cotesia glomerata* female parasitoid wasps (n = 12); Pb, *Pieris brassicae* caterpillars (n = 10); Pb-Cg, *P. brassicae* caterpillars in the presence of *C. glomerata* female parasitoid wasps (n = 14); Pb-Fr, *P. brassicae* caterpillar frass (n = 10); and Pb-S, *P. brassicae* spit (n = 12), are listed. Relative amounts of volatiles are presented as average peak height (SE)/$10^4$. The volatiles are listed according to their elution order in a chromatographic window. Volatiles with Variable Importance in the Projection (VIP) scores equal to or higher than 1.0 (presented in bold), are considered important in separating the different treatment groups of the given analysis. Significant differences among the five treatments detected by Kruskal-Wallis test with Dunn's post-hoc test for multiple comparisons are indicated by letters in the table. Different letters indicate significant difference (P < 0.05). NF, not found.

| Number | Compound and class | Cg | Pb | Pb-Cg | Pb-S | Pb-Fr | VIP-SCORE |
|---|---|---|---|---|---|---|---|
| 1 | 2,3-Butanedione | 9.9 (1.2) a | 115.9 (17.7) b | 290.9 (106.5) b | 167.1 (33.1) b | 1540.7 (712.9) b | **1.04** |
| 2 | 2-Butenenitrile | NF a | NF a | 1.2 (0.6) a | 11.3 (3) b | NF a | **1.17** |
| 3 | 3-Methylbutanal | 0.1 (0.1) a | 2 (0.4) ab | 3.4 (0.5) bc | 243.4 (21.1) d | 92.1 (31.5) cd | 0.91 |
| 4 | 2-Methylbutanal | 0.1 (0.1) a | 1.6 (0.3) ab | 3.2 (0.5) bc | 277.9 (23.7) d | 108.1 (36.6) cd | 0.91 |
| 5 | 1-Methoxy-2-propanol | 3.1 (1.3) a | 74.8 (15.3) b | 256.5 (93.6) b | 149.2 (9.6) b | 538.2 (259.6) b | **1.04** |
| 6 | 1-Penten-3-ol | NF a | 55 (11.9) ab | 158.7 (51.8) b | 466.6 (64.3) c | 1028.1 (414) bc | 0.98 |
| 7 | Pentanal | 7.2 (0.8) a | 8.4 (1.1) a | 9.6 (2.6) a | 358.6 (29.8) b | 22.9 (12) a | **1.11** |
| 8 | Methyl thiocyanate | NF a | NF a | NF a | 69.5 (7.9) b | 86.6 (31.9) b | 0.98 |
| 9 | 3-Methyl-1-butanol | NF a | NF a | 1.6 (1.2) a | 82.9 (39.7) b | 90.4 (44.3) b | 0.78 |
| 10 | 3-Penten-2-one | 1.2 (0.6) a | 2.6 (1.1) a | 4.5 (1.6) a | 104.3 (26.6) b | 122 (50.3) b | 0.82 |
| 11 | Unknown | NF a | 0.2 (0.2) a | 0.3 (0.2) a | 62.8 (5.7) b | 324.4 (162.9) ab | 0.85 |
| 12 | Dimethyl disulfide | 1.5 (0.6) a | 182.5 (23.9) ab | 780.2 (131.3) bc | 2738 (296.5) c | 3182.7 (589) c | **1.10** |
| 13 | (Z)-2-Pentenal | NF a | 1.4 (0.6) a | 3.1 (0.8) ab | 102.2 (7.5) c | 36.4 (14.8) bc | 0.86 |
| 14 | 3-Methyl-3-butenenitrile | NF a | NF a | 1 (0.4) a | 12.4 (2.7) b | 0.2 (0.1) a | **1.30** |
| 15 | Methylthioacetaldehyde | NF a | NF a | NF a | 0.1 (0.1) a | 90.9 (32.1) b | **1.40** |
| 16 | 1-Pentanol | 1.7 (0.9) a | 1.5 (0.8) a | 3.7 (1.5) ab | 220.8 (56.8) c | 58.1 (23.1) bc | 0.81 |
| 17 | (Z)-2-Penten-1-ol | NF a | NF a | 0.6 (0.4) a | 63.4 (7.8) b | 40.1 (18) b | 0.87 |
| 18 | 2,3-Butanediol | NF a | 1.1 (1.1) a | 19.5 (10.6) ab | 23.4 (5.4) b | 187.8 (94.9) ab | 0.93 |
| 19 | Ethyl methanesulfinate | NF a | NF a | NF a | 31.4 (8.8) b | 11.3 (3.7) b | 0.95 |
| 20 | 3-Methylbutanoic acid | NF a | NF a | 0.3 (0.1) a | 18.7 (1.8) b | 3.1 (1.1) ab | 0.89 |
| 21 | 2,3-Heptanedione | NF a | NF a | NF a | 4.6 (0.5) b | NF a | **1.14** |
| 22 | (Z)-3-Hexen-1-ol | 2.3 (1.5) a | 42.2 (5.3) ab | 98.7 (23.4) b | 3384.9 (454.5) c | 1332.7 (569.4) bc | 0.97 |
| 23 | (Z)-2-Hexen-1-ol | NF a | NF a | 0.6 (0.5) a | 99.8 (22.7) b | 15.5 (7.6) a | 0.94 |
| 24 | 1-Hexanol | 0.3 (0.3) a | NF a | 2.3 (1.3) a | 70.8 (21.6) b | 43.5 (17.5) b | 0.86 |
| 25 | Cyclohexanol | NF a | 9 (2.8) b | 25.4 (7.1) b | 4 (1.2) ab | 397.9 (197.6) b | **1.06** |
| 26 | 3-Ethyl-1,5-octadiene, Isomer I | NF a | 13.3 (1.8) b | 10.9 (1.8) b | 0.2 (0.2) a | NF a | **1.15** |
| 27 | 3-Ethyl-1,5-octadiene, Isomer II | NF a | 53.7 (7.3) b | 48.5 (6.5) b | NF a | NF a | **1.18** |
| 28 | 3-(Methylthio)propanal | NF a | NF a | NF a | 24.5 (3.3) b | 6.2 (3.2) a | 0.99 |
| 29 | Unknown | NF a | NF a | NF a | 42 (7.4) b | 3.6 (2.4) a | 0.97 |
| 30 | 3,7-Decadiene, Isomer I | NF a | 9.9 (0.8) bc | 12.2 (2.6) bc | 19.8 (3.9) c | 2.8 (1.1) ab | **1.20** |
| 31 | Dimethyl trisulfide | NF a | 19 (2.9) ab | 249.4 (48.4) bc | 6948 (557.9) d | 2099.9 (354.5) cd | **1.06** |
| 32 | 3,7-Decadiene, Isomer II | NF a | 6.3 (1.3) bc | 9.9 (2.4) c | 23 (5.2) c | 1.2 (0.9) ab | **1.04** |
| 33 | (E,E)-2,4-Heptadienal | NF a | NF a | NF b | 5.9 (1.1) b | 5.8 (2.2) b | 0.86 |
| 34 | Phenylacetaldehyde | 5.1 (0.4) a | 5 (0.5) a | 6.8 (1) a | 38.5 (3.5) b | 11.6 (3) a | 0.94 |
| 35 | 3-Methyl-2-butenyl 2-methylbutanoate | 0.8 (0.5) a | NF a | NF a | 140.3 (8.5) b | 56.5 (23.9) b | 0.94 |
| 36 | Methyl (methylthio)methyl disulfide | NF a | 0.2 (0.1) a | 6.7 (2) b | 76.3 (7.9) c | 72.5 (28.6) bc | **1.19** |
| 37 | Benzyl cyanide | 1.3 (0.3) a | 2.8 (0.6) ab | 10.7 (2.2) bc | 549.6 (66.4) d | 31 (11) cd | 0.95 |
| 38 | 4-Ketoisophorone | NF a | NF a | NF a | 3.9 (0.3) b | 7.7 (3.4) b | 0.94 |
| 39 | beta-Cyclocitral | NF a | 0.4 (0.1) a | 0.6 (0.2) a | 28.2 (1.4) b | 77.8 (28.7) b | 0.93 |
| 40 | Dimethyl tetrasulfide | NF a | NF a | 0.6 (0.2) a | 2119.4 (226.8) b | 145.5 (36) b | 0.90 |
| 41 | Chavibetol | NF a | 0.3 (0.2) a | 1.9 (1) a | 30.1 (3.5) b | 77.6 (37.2) b | 0.87 |
| 42 | (E)-beta-Ionone | NF a | 0.5 (0.5) a | 3 (1.1) a | 104.8 (5) b | 145.9 (53.5) b | 0.91 |
| 43 | beta-Ionone epoxide | NF a | 0.5 (0.3) a | 1.3 (0.4) a | 51 (3.4) b | 44.6 (17.8) b | 0.89 |
| 44 | Dihydroactinidiolide | NF a | 0.1 (0.1) a | 0.2 (0.1) a | 4.7 (0.5) b | 3.4 (1.4) b | 0.79 |
| 45 | Tricyclopentadeca-3,7-dien[8.4.0.1(11,14)] | 0.7 (0.2) a | 10.9 (0.7) b | 8.8 (0.9) b | 3.4 (0.2) a | 1.6 (0.6) a | **1.20** |

**Table S4. Volatiles used in separating *Cotesia glomerata* female wasps (Cg, n = 12), *Pieris brassicae* caterpillars (Pb, n = 10) and the interaction of *P. brassicae* caterpillars with *C. glomerata* female wasps (Pb-Cg, n = 14) sample treatments (Fig S5).** Volatiles are listed according to ranking order of their Variable Importance in the Projection (VIP) scores values, where those with VIP scores of equal to or higher than 1.0, are considered important in separating the treatment groups of the given analysis.

| Primary ID | Chemical name | VIP-Score |
|---|---|---|
| 36 | Methyl (methylthio)methyl disulfide | 1.56332 |
| 31 | Dimethyl trisulfide | 1.39172 |
| 14 | 3-Methyl-3-butenenitrile | 1.37969 |
| 27 | 3-Ethyl-1,5-octadiene, Isomer II | 1.37419 |
| 30 | 3,7-Decadiene, Isomer I | 1.35815 |
| 26 | 3-Ethyl-1,5-octadiene, Isomer I | 1.34633 |
| 12 | Dimethyl disulfide | 1.31333 |
| 6 | 1-Penten-3-ol | 1.29147 |
| 25 | Cyclohexanol | 1.26832 |
| 32 | 3,7-Decadiene, Isomer II | 1.26563 |
| 45 | Tricyclopentadeca-3,7-dien[8.4.0.1(11,14)] | 1.26062 |
| 3 | 3-Methylbutanal | 1.21712 |
| 4 | 2-Methylbutanal | 1.20836 |
| 22 | (Z)-3-Hexen-1-ol | 1.20629 |
| 5 | 1-Methoxy-2-propanol | 1.19415 |
| 1 | 2,3-Butanedione | 1.1492 |
| 40 | Dimethyl tetrasulfide | 1.09912 |
| 37 | Benzyl cyanide | 1.09685 |
| 24 | 1-Hexanol | 1.06427 |
| 2 | 2-Butenenitrile | 1.06105 |
| 13 | (Z)-2-Pentenal | 0.978154 |
| 11 | Unknown | 0.974729 |
| 17 | (Z)-2-Penten-1-ol | 0.965661 |
| 39 | beta-Cyclocitral | 0.942238 |
| 43 | beta-Ionone epoxide | 0.933374 |
| 9 | 3-Methyl-1-butanol | 0.932564 |
| 41 | Chavibetol | 0.928908 |
| 34 | Phenylacetaldehyde | 0.832774 |
| 18 | 2,3-Butanediol | 0.826549 |
| 42 | (E)-beta-Ionone | 0.809716 |
| 33 | (E,E)-2,4-Heptadienal | 0.705911 |
| 20 | 3-Methylbutanoic acid | 0.663228 |
| 28 | 3-(Methylthio)propanal | 0.652614 |
| 23 | (Z)-2-Hexen-1-ol | 0.635438 |
| 21 | 2,3-Heptanedione | 0.633611 |
| 38 | 4-Ketoisophorone | 0.62339 |
| 7 | Pentanal | 0.575643 |
| 10 | 3-Penten-2-one | 0.557406 |
| 35 | 3-Methyl-2-butenyl 2-methylbutanoate | 0.554326 |
| 8 | Methyl thiocyanate | 0.526719 |
| 19 | Ethyl methanesulfinate | 0.514083 |
| 16 | 1-Pentanol | 0.505486 |
| 44 | Dihydroactinidiolide | 0.502176 |
| 15 | Methylthioacetaldehyde | 0.476651 |
| 29 | Unknown | 0.132699 |

**Table S5. Volatiles used in separating *Pieris brassicae* caterpillars (Pb, n = 10) and the interaction of *P. brassicae* caterpillars with *C. glomerata* female wasps (Pb-Cg, n = 14) sample treatments (Fig S6).** Volatiles are listed according to ranking order of their Variable Importance in the Projection (VIP) scores values, where those with VIP scores of equal to or higher than 1.0, are considered important in separating the treatment groups of the given analysis.

| Primary ID | Chemical name | VIP-Score |
|---|---|---|
| 36 | Methyl (methylthio)methyl disulfide | 2.07587 |
| 12 | Dimethyl disulfide | 1.92275 |
| 31 | Dimethyl trisulfide | 1.69443 |
| 14 | 3-Methyl-3-butenenitrile | 1.5985 |
| 37 | Benzyl cyanide | 1.51645 |
| 24 | 1-Hexanol | 1.27124 |
| 35 | 3-Methyl-2-butenyl 2-methylbutanoate | 1.24411 |
| 40 | Dimethyl tetrasulfide | 1.23794 |
| 2 | 2-Butenenitrile | 1.18014 |
| 45 | Tricyclopentadeca-3,7-dien[8.4.0.1(11,14)] | 1.16026 |
| 11 | Unknown | 1.11114 |
| 17 | (Z)-2-Penten-1-ol | 1.06676 |
| 9 | 3-Methyl-1-butanol | 1.03313 |
| 41 | Chavibetol | 0.999371 |
| 43 | beta-Ionone epoxide | 0.98264 |
| 18 | 2,3-Butanediol | 0.957447 |
| 34 | Phenylacetaldehyde | 0.942159 |
| 13 | (Z)-2-Pentenal | 0.916875 |
| 28 | 3-(Methylthio)propanal | 0.909736 |
| 39 | beta-Cyclocitral | 0.90317 |
| 6 | 1-Penten-3-ol | 0.890276 |
| 4 | 2-Methylbutanal | 0.877197 |
| 22 | (Z)-3-Hexen-1-ol | 0.857471 |
| 42 | (E)-beta-Ionone | 0.851761 |
| 26 | 3-Ethyl-1,5-octadiene, Isomer I | 0.842136 |
| 1 | 2,3-Butanedione | 0.839521 |
| 3 | 3-Methylbutanal | 0.831809 |
| 5 | 1-Methoxy-2-propanol | 0.81872 |
| 25 | Cyclohexanol | 0.785347 |
| 30 | 3,7-Decadiene, Isomer I | 0.783212 |
| 38 | 4-Ketoisophorone | 0.746923 |
| 33 | (E,E)-2,4-Heptadienal | 0.728538 |
| 21 | 2,3-Heptanedione | 0.71051 |
| 23 | (Z)-2-Hexen-1-ol | 0.710482 |
| 20 | 3-Methylbutanoic acid | 0.683183 |
| 27 | 3-Ethyl-1,5-octadiene, Isomer II | 0.642632 |
| 7 | Pentanal | 0.631077 |
| 16 | 1-Pentanol | 0.605405 |
| 32 | 3,7-Decadiene, Isomer II | 0.590844 |
| 19 | Ethyl methanesulfinate | 0.557555 |
| 8 | Methyl thiocyanate | 0.55321 |
| 10 | 3-Penten-2-one | 0.465652 |
| 29 | Unknown | 0.322982 |
| 44 | Dihydroactinidiolide | 0.273144 |
| 15 | Methylthioacetaldehyde | 0.0785777 |

**Table S6. Volatiles used in separating the interaction of *Pieris brassicae* caterpillars with *Cotesia glomerata* female wasps (Pb-Cg, n = 14) and *P. brassicae* caterpillar spit (Pb-S, n = 12), sample treatments (Fig S7).** Volatiles are listed according to ranking order of their Variable Importance in the Projection (VIP) scores values, where those with VIP scores of equal to or higher than 1.0, are considered important in separating the treatment groups of the given analysis.

| Primary ID | Chemical name | VIP-Score |
|---|---|---|
| 35 | 3-Methyl-2-butenyl 2-methylbutanoate | 1.18987 |
| 8 | Methyl thiocyanate | 1.18927 |
| 38 | 4-Ketoisophorone | 1.18925 |
| 21 | 2,3-Heptanedione | 1.18687 |
| 28 | 3-(Methylthio)propanal | 1.18366 |
| 40 | Dimethyl tetrasulfide | 1.16523 |
| 37 | Benzyl cyanide | 1.164 |
| 11 | Unknown | 1.15829 |
| 31 | Dimethyl trisulfide | 1.15172 |
| 23 | (Z)-2-Hexen-1-ol | 1.13638 |
| 17 | (Z)-2-Penten-1-ol | 1.13498 |
| 20 | 3-Methylbutanoic acid | 1.12993 |
| 34 | Phenylacetaldehyde | 1.12219 |
| 4 | 2-Methylbutanal | 1.11885 |
| 39 | beta-Cyclocitral | 1.11868 |
| 19 | Ethyl methanesulfinate | 1.11794 |
| 3 | 3-Methylbutanal | 1.1088 |
| 43 | beta-Ionone epoxide | 1.07481 |
| 16 | 1-Pentanol | 1.05818 |
| 42 | (E)-beta-Ionone | 1.05223 |
| 29 | Unknown | 1.04776 |
| 33 | (E,E)-2,4-Heptadienal | 1.04304 |
| 27 | 3-Ethyl-1,5-octadiene, Isomer II | 1.03943 |
| 41 | Chavibetol | 1.03686 |
| 13 | (Z)-2-Pentenal | 1.03618 |
| 36 | Methyl (methylthio)methyl disulfide | 1.01861 |
| 12 | Dimethyl disulfide | 1.00794 |
| 22 | (Z)-3-Hexen-1-ol | 1.00726 |
| 44 | Dihydroactinidiolide | 1.0004 |
| 45 | Tricyclopentadeca-3,7-dien[8.4.0.1(11,14)] | 0.996387 |
| 7 | Pentanal | 0.994438 |
| 14 | 3-Methyl-3-butenenitrile | 0.974921 |
| 10 | 3-Penten-2-one | 0.971786 |
| 26 | 3-Ethyl-1,5-octadiene, Isomer I | 0.935729 |
| 24 | 1-Hexanol | 0.901818 |
| 6 | 1-Penten-3-ol | 0.856495 |
| 2 | 2-Butenenitrile | 0.855662 |
| 9 | 3-Methyl-1-butanol | 0.795921 |
| 25 | Cyclohexanol | 0.680023 |
| 1 | 2,3-Butanedione | 0.612927 |
| 5 | 1-Methoxy-2-propanol | 0.589883 |
| 18 | 2,3-Butanediol | 0.513856 |
| 30 | 3,7-Decadiene, Isomer I | 0.490609 |
| 32 | 3,7-Decadiene, Isomer II | 0.399193 |
| 15 | Methylthioacetaldehyde | 0.293689 |

**Table S7. Volatiles used in separating the *Pieris brassicae* caterpillar spit (Pb-S, n = 12), and *P. brassicae* caterpillar frass (Pb-Fr, n = 10) sample treatments (Fig S8).** Volatiles are listed according to ranking order of their Variable Importance in the Projection (VIP) scores values, where those with VIP scores of equal to or higher than 1.0, are considered important in separating the treatment groups of the given analysis.

| Primary ID | Chemical name | VIP-Score |
|---|---|---|
| 21 | 2,3-Heptanedione | 1.47069 |
| 14 | 3-Methyl-3-butenenitrile | 1.41146 |
| 37 | Benzyl cyanide | 1.37845 |
| 15 | Methylthioacetaldehyde | 1.3601 |
| 34 | Phenylacetaldehyde | 1.26864 |
| 2 | 2-Butenenitrile | 1.26709 |
| 40 | Dimethyl tetrasulfide | 1.22517 |
| 28 | 3-(Methylthio)propanal | 1.2005 |
| 13 | (Z)-2-Pentenal | 1.19752 |
| 23 | (Z)-2-Hexen-1-ol | 1.19502 |
| 20 | 3-Methylbutanoic acid | 1.17384 |
| 3 | 3-Methylbutanal | 1.17326 |
| 4 | 2-Methylbutanal | 1.15938 |
| 7 | Pentanal | 1.15138 |
| 31 | Dimethyl trisulfide | 1.15035 |
| 22 | (Z)-3-Hexen-1-ol | 1.13272 |
| 45 | Tricyclopentadeca-3,7-dien[8.4.0.1(11,14)] | 1.11074 |
| 30 | 3,7-Decadiene, Isomer I | 1.09918 |
| 29 | Unknown | 1.08182 |
| 32 | 3,7-Decadiene, Isomer II | 1.04475 |
| 17 | (Z)-2-Penten-1-ol | 1.03121 |
| 16 | 1-Pentanol | 1.0235 |
| 35 | 3-Methyl-2-butenyl 2-methylbutanoate | 1.01617 |
| 43 | beta-Ionone epoxide | 0.971688 |
| 11 | Unknown | 0.952159 |
| 36 | Methyl (methylthio)methyl disulfide | 0.924777 |
| 38 | 4-Ketoisophorone | 0.854173 |
| 25 | Cyclohexanol | 0.853377 |
| 1 | 2,3-Butanedione | 0.847814 |
| 39 | beta-Cyclocitral | 0.843729 |
| 8 | Methyl thiocyanate | 0.841366 |
| 42 | (E)-beta-Ionone | 0.82494 |
| 41 | Chavibetol | 0.822314 |
| 5 | 1-Methoxy-2-propanol | 0.803075 |
| 10 | 3-Penten-2-one | 0.765907 |
| 19 | Ethyl methanesulfinate | 0.764512 |
| 33 | (E,E)-2,4-Heptadienal | 0.702099 |
| 6 | 1-Penten-3-ol | 0.647433 |
| 12 | Dimethyl disulfide | 0.610209 |
| 9 | 3-Methyl-1-butanol | 0.607502 |
| 18 | 2,3-Butanediol | 0.570747 |
| 24 | 1-Hexanol | 0.567647 |
| 44 | Dihydroactinidiolide | 0.52246 |
| 26 | 3-Ethyl-1,5-octadiene, Isomer I | 0.333459 |
| 27 | 3-Ethyl-1,5-octadiene, Isomer II | 0.191552 |

**Table S8. Volatiles used in separating the *Pieris brassicae* caterpillars (Pb, n = 10), the interaction of *P. brassicae* caterpillars with *Cotesia glomerata* female wasps (Pb-Cg, n = 14), and *P. brassicae* caterpillar frass (Pb-Fr, n = 10) sample treatments (Fig S9).** Volatiles are listed according to ranking order of their Variable Importance in the Projection (VIP) scores values, where those with VIP scores of equal to or higher than 1.0, are considered important in separating the treatment groups of the given analysis.

| Primary ID | Chemical name | VIP-Score |
|---|---|---|
| 14 | 3-Methyl-3-butenenitrile | 1.53713 |
| 36 | Methyl (methylthio)methyl disulfide | 1.49888 |
| 37 | Benzyl cyanide | 1.44649 |
| 31 | Dimethyl trisulfide | 1.37488 |
| 12 | Dimethyl disulfide | 1.27985 |
| 8 | Methyl thiocyanate | 1.16994 |
| 15 | Methylthioacetaldehyde | 1.16173 |
| 35 | 3-Methyl-2-butenyl 2-methylbutanoate | 1.12286 |
| 19 | Ethyl methanesulfinate | 1.12207 |
| 38 | 4-Ketoisophorone | 1.09766 |
| 40 | Dimethyl tetrasulfide | 1.08304 |
| 2 | 2-Butenenitrile | 1.0747 |
| 21 | 2,3-Heptanedione | 1.06918 |
| 39 | beta-Cyclocitral | 1.032 |
| 27 | 3-Ethyl-1,5-octadiene, Isomer II | 1.03194 |
| 33 | (E,E)-2,4-Heptadienal | 0.999281 |
| 42 | (E)-beta-Ionone | 0.991173 |
| 26 | 3-Ethyl-1,5-octadiene, Isomer I | 0.989933 |
| 24 | 1-Hexanol | 0.984174 |
| 45 | Tricyclopentadeca-3,7-dien[8.4.0.1(11,14)] | 0.970013 |
| 4 | 2-Methylbutanal | 0.965192 |
| 17 | (Z)-2-Penten-1-ol | 0.958219 |
| 43 | beta-Ionone epoxide | 0.958126 |
| 3 | 3-Methylbutanal | 0.940685 |
| 41 | Chavibetol | 0.93559 |
| 9 | 3-Methyl-1-butanol | 0.927445 |
| 10 | 3-Penten-2-one | 0.892843 |
| 20 | 3-Methylbutanoic acid | 0.889393 |
| 22 | (Z)-3-Hexen-1-ol | 0.888146 |
| 18 | 2,3-Butanediol | 0.880791 |
| 11 | Unknown | 0.866956 |
| 28 | 3-(Methylthio)propanal | 0.852439 |
| 13 | (Z)-2-Pentenal | 0.84884 |
| 32 | 3,7-Decadiene, Isomer II | 0.840748 |
| 44 | Dihydroactinidiolide | 0.821653 |
| 23 | (Z)-2-Hexen-1-ol | 0.818063 |
| 1 | 2,3-Butanedione | 0.807042 |
| 5 | 1-Methoxy-2-propanol | 0.78006 |
| 34 | Phenylacetaldehyde | 0.774451 |
| 7 | Pentanal | 0.767969 |
| 16 | 1-Pentanol | 0.766034 |
| 30 | 3,7-Decadiene, Isomer I | 0.764335 |
| 25 | Cyclohexanol | 0.72219 |
| 29 | Unknown | 0.695 |
| 6 | 1-Penten-3-ol | 0.662903 |

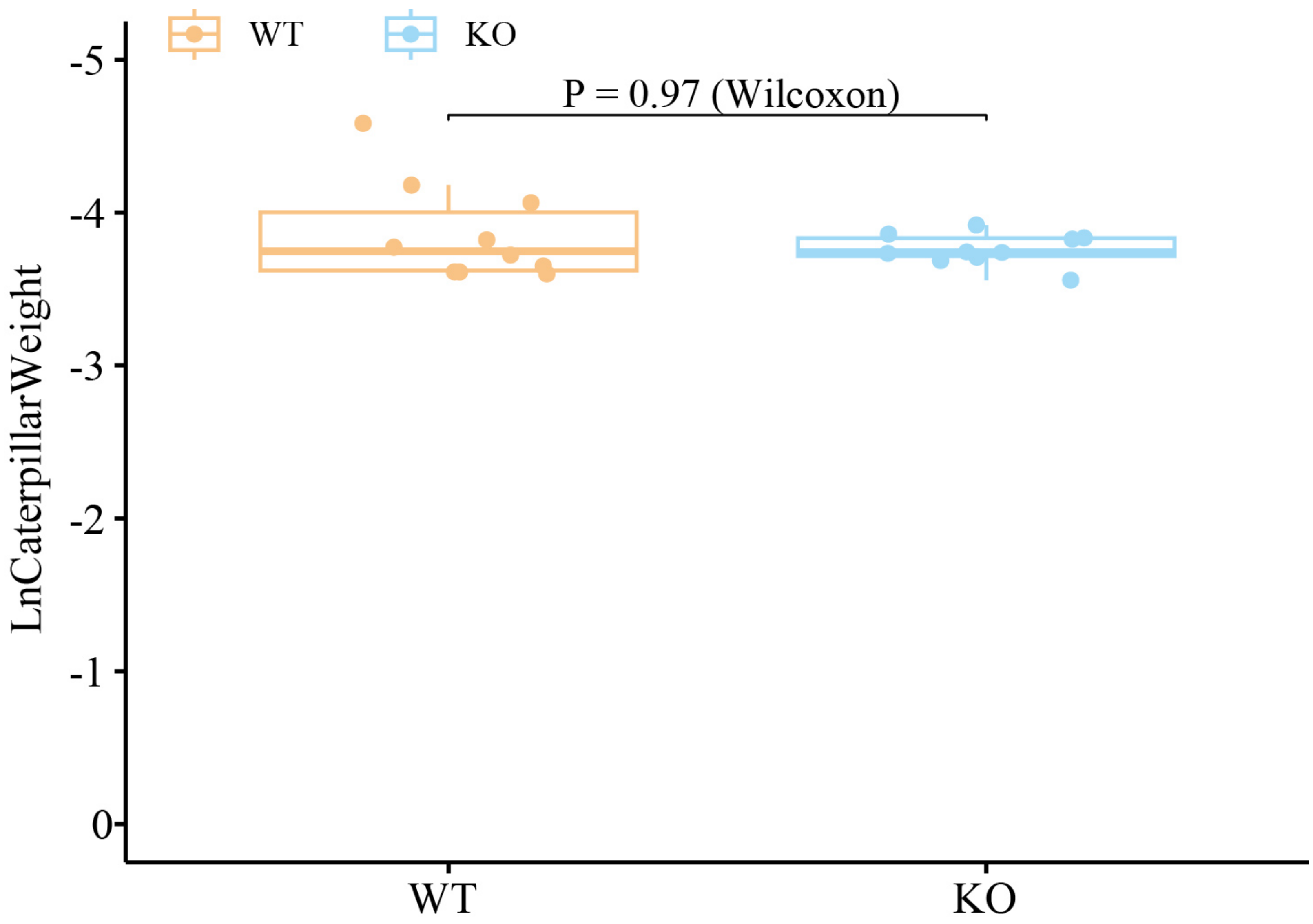


**Fig S1.** ***Pieris brassicae*** **caterpillar development in a Petri dish environment.** X axis indicates the date since caterpillars have been deposited in the Petri dish. Y axis indicates the weight of caterpillars (Ln Weight). Caterpillar weight (n = 10 for both genotypes) in this figure is presented by the mean of a group of caterpillars weight in each Petri dish on day 6.

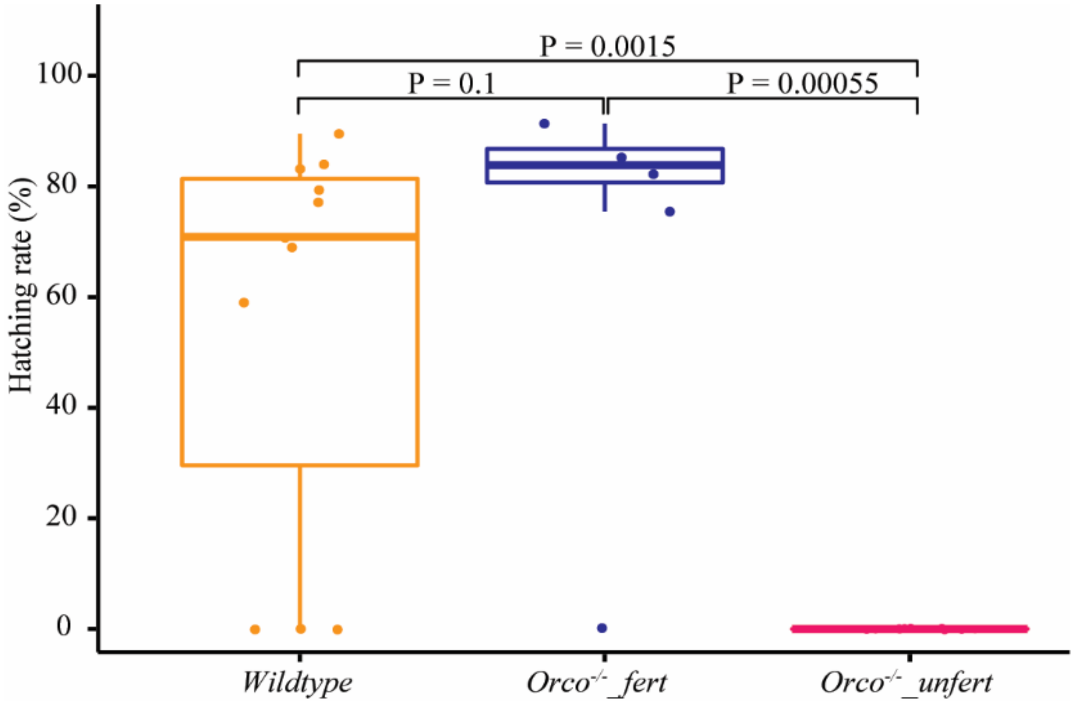


**Fig S2. Egg-hatching rates by *wildtype* (n=11), *Orco* $^{-/-}$ *_fert* (n=5) and *Orco* $^{-/-}$ *_unfert* (n=9) butterflies.** Orco-/-_fert represents mated Orco female butterflies, Orco-/-_unfert represents unmated female butterflies. Differences were tested by using Student's t-test.

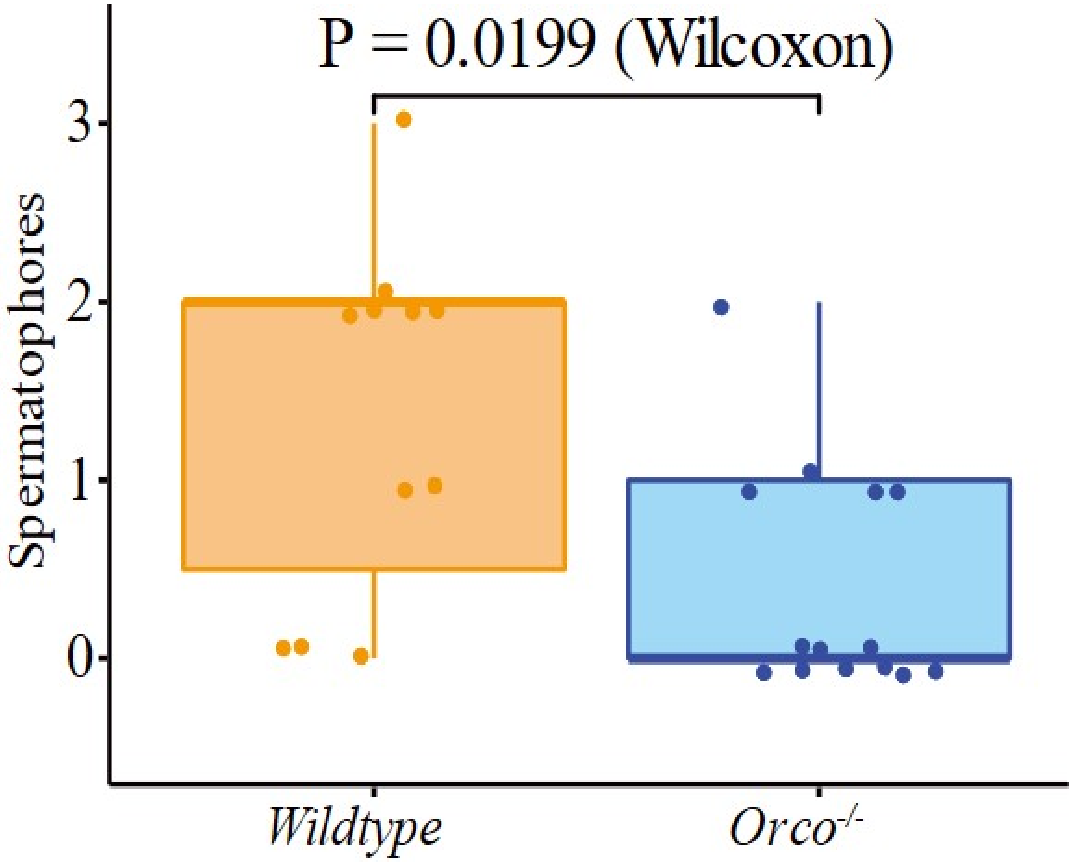


**Fig S3. Mating frequency of *Pieris brassicae* butterflies.** The boxplots indicate the number of spermatophores in WT female butterflies (n = 11) and in KO butterflies (*Orco* $^{-/-}$) (n = 14), the difference was tested by using Wilcoxon rank-sum test. WT and Orco KO butterfly spermatophores are indicated in orange and blue respectively.

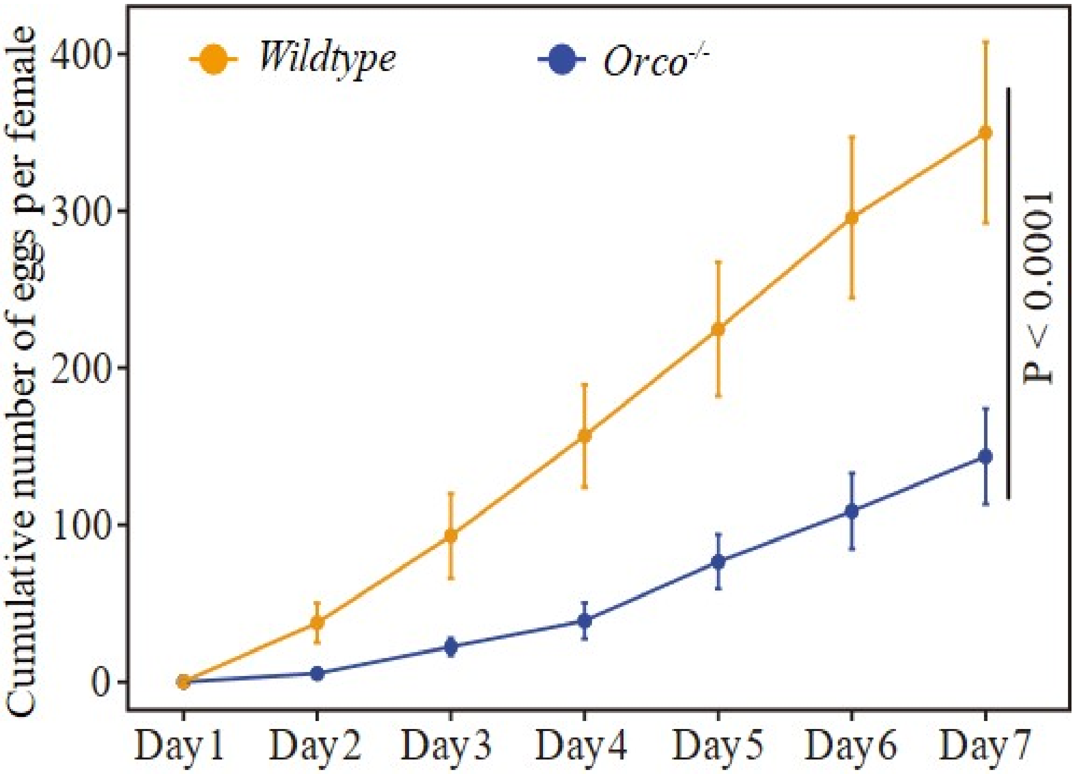


**Fig S4. *Pieris brassicae wildtype* and *Orco* knockout (*Orco* $^{-/-}$) butterfly oviposition dynamics.** The orange line indicates the number of eggs per female (mean ± SE) laid by wildtype (WT) butterflies (n = 11), the blue line indicates the number of eggs laid by *Orco* knockout (KO) butterflies (n = 14). Differences were analyzed using a GLM with negative binominal distribution, difference test results are shown in the line chart.

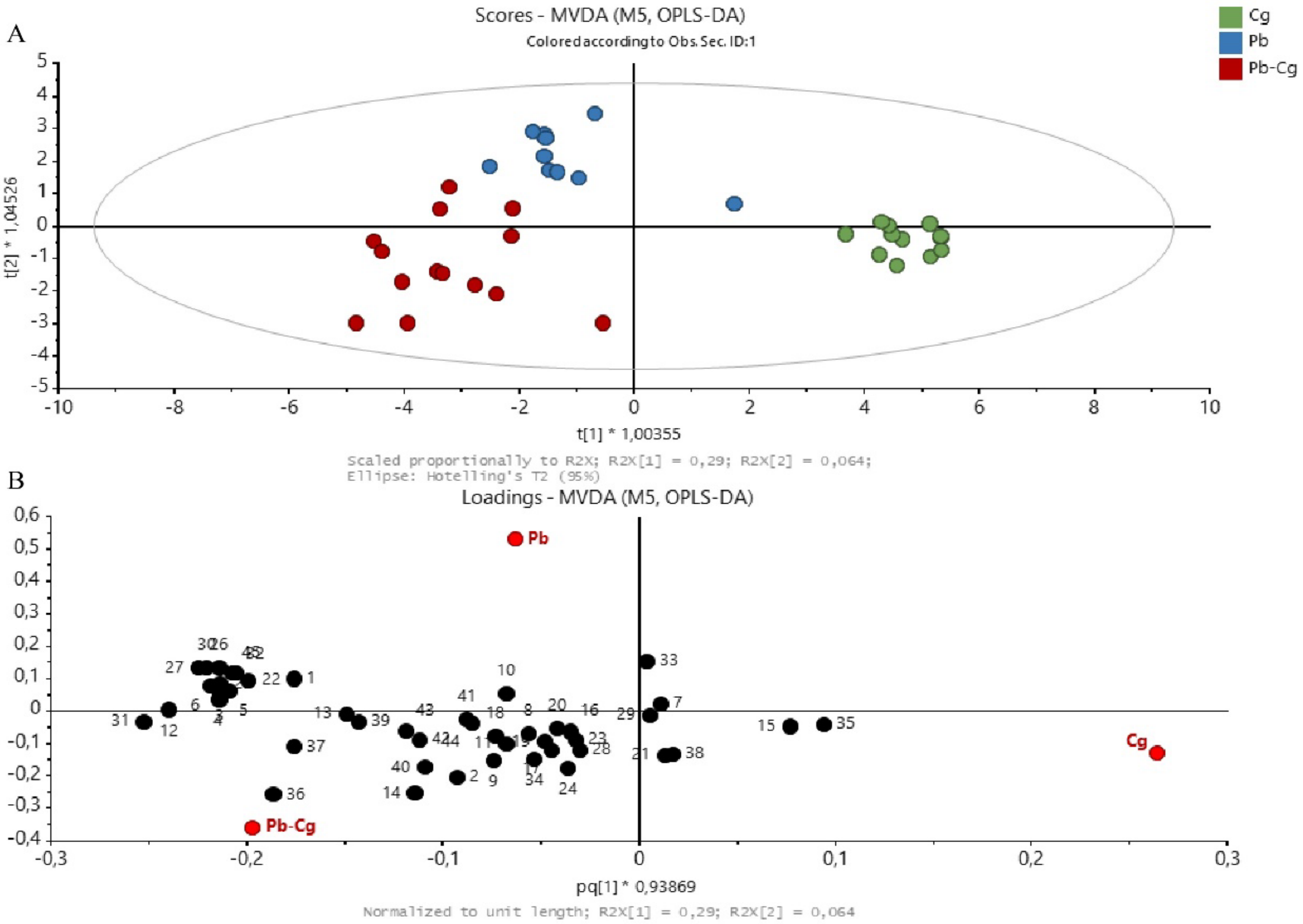


**Fig S5. Overview of volatile blends of *Pieris brassicae* caterpillars, *Cotesia glomerata* parasitoid wasps and caterpillars-parasitoid in interaction.** (A), OPLS-DA (Orthogonal Projection to Latent Structures Discriminant Analysis) two-dimensional score plot of treatment groups: Cg, *Cotesia glomerata* female parasitoid wasps (n = 12), based on their volatile content; Pb, *Pieris brassicae* caterpillars (n = 10); Pb-Cg, *P. brassicae* caterpillars in the presence of *C. glomerata* female parasitoid wasps (n = 14. (B), Loading plot showing the contribution of each identified volatile compound to the separation of the different treatments. Volatiles closer to the treatment in the plot indicate higher correlation. Numbers in the loading plot refer to the volatile compounds listed in Supporting information Table S3.

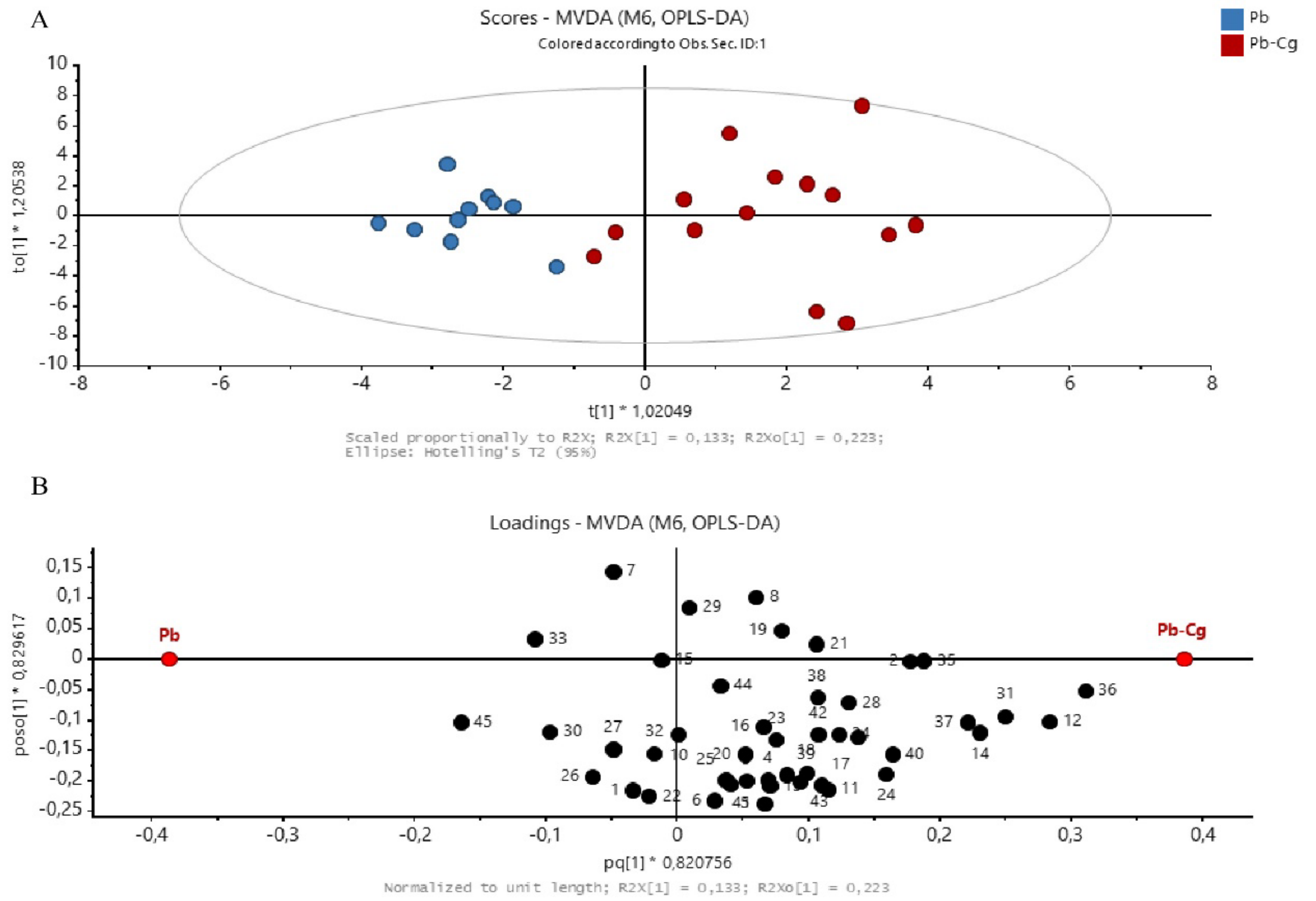


**Fig S6. Overview of volatile blends of *Pieris brassicae* caterpillars and caterpillars-parasitoid (*Cotesia glomerata*) in interaction.** (A), OPLS-DA (Orthogonal Projection to Latent Structures Discriminant Analysis) two-dimensional score plot of treatment groups: Pb, *Pieris brassicae* caterpillars (n = 10); Pb-Cg, *P. brassicae* caterpillars in the presence of *C. glomerata* female parasitoid wasps (n = 14), based on their volatile content. (B), Loading plot showing the contribution of each identified volatile compound to the separation of the different treatments. Volatiles closer to the treatment in the plot indicate higher correlation. Numbers in the loading plot refer to the volatile compounds listed in Supporting information Table S3.

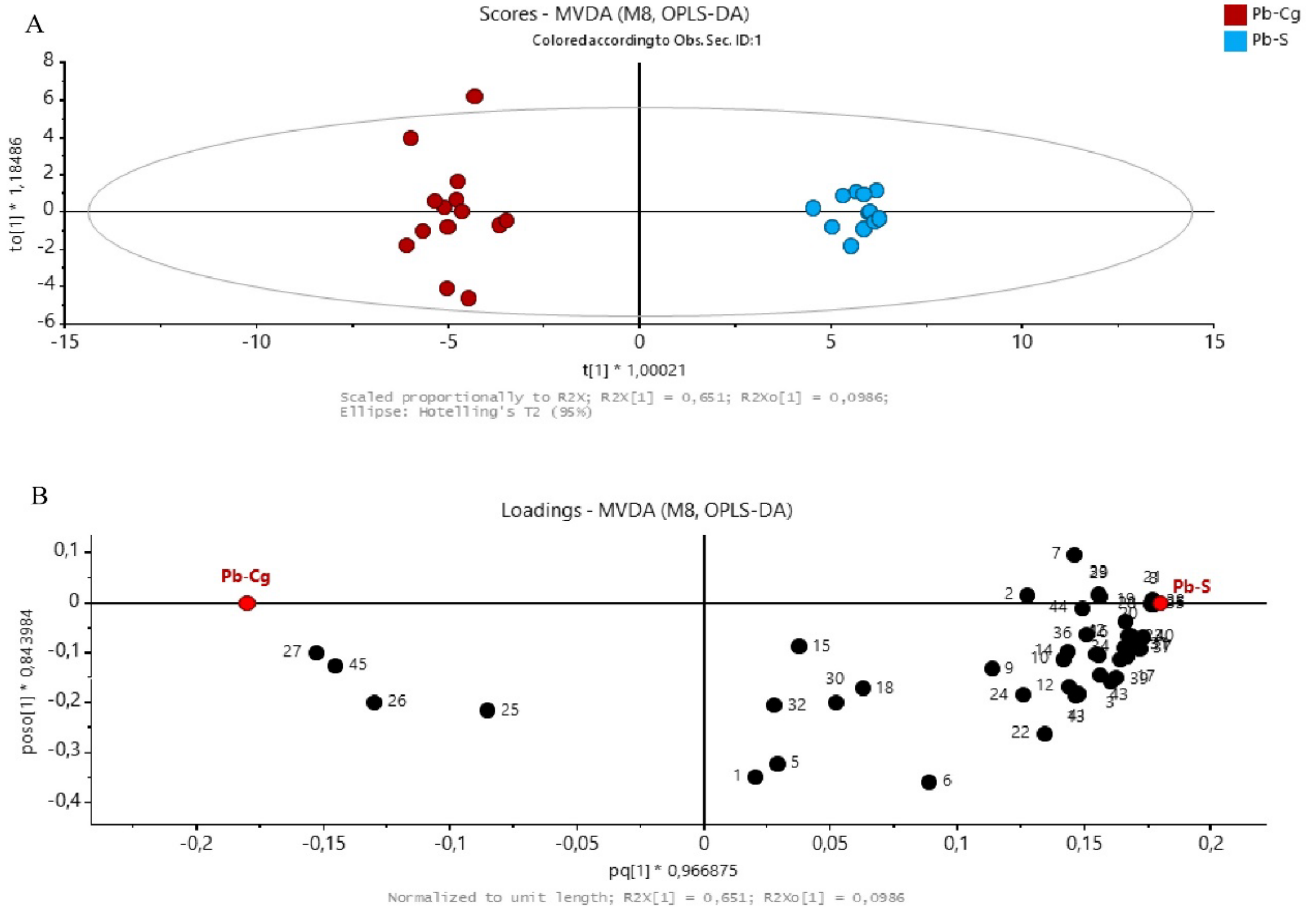


**Fig S7. Overview of volatile blends of *Pieris brassicae* caterpillar spit and caterpillars-parasitoid wasp (*Cotesia glomerata*) in interaction.** (A), OPLS-DA (Orthogonal Projection to Latent Structures Discriminant Analysis) two-dimensional score plot of treatment groups: *P. brassicae* spit (n = 12), Pb-Cg, *P. brassicae* caterpillars in the presence of *C. glomerata* female parasitoid wasps (n = 14), based on their volatile content. (B), Loading plot showing the contribution of each identified volatile compound to the separation of the different treatments. Volatiles closer to the treatment in the plot indicate higher correlation. Numbers in the loading plot refer to the volatile compounds listed in Supporting information Table S3.

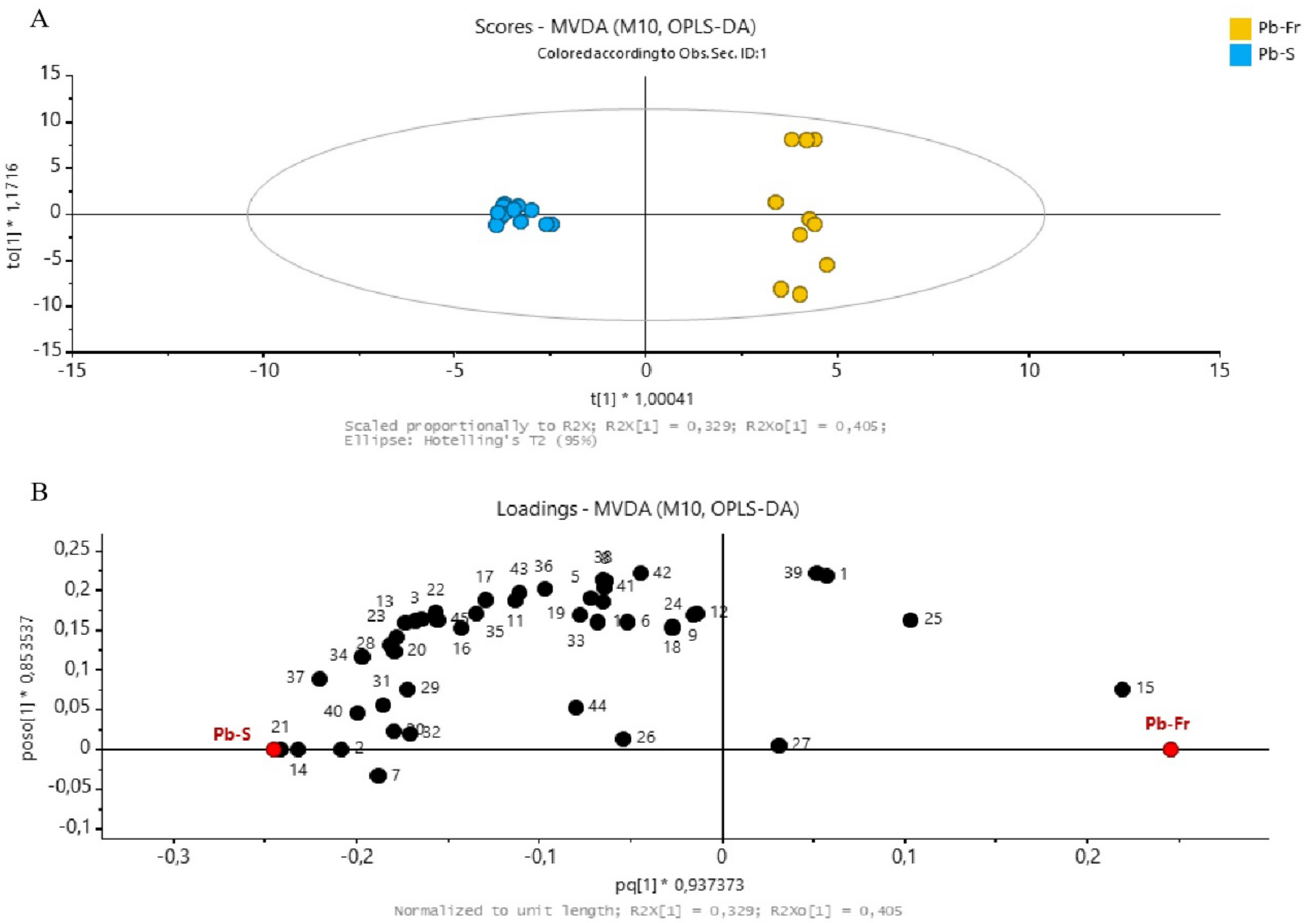


**Fig S8. Overview of volatile blends of *Pieris brassicae* caterpillar spit and *P. brassicae* caterpillar frass.** (A), OPLS-DA (Orthogonal Projection to Latent Structures Discriminant Analysis) two-dimensional score plot of treatment groups: Pb-Fr, *P. brassicae* caterpillar frass (n = 10); and Pb-S, *P. brassicae* spit (n = 12), based on their volatile content. (B), Loading plot showing the contribution of each identified volatile compound to the separation of the different treatments. Volatiles closer to the treatment in the plot indicate higher correlation. Numbers in the loading plot refer to the volatile compounds listed in Supporting information Table S3.

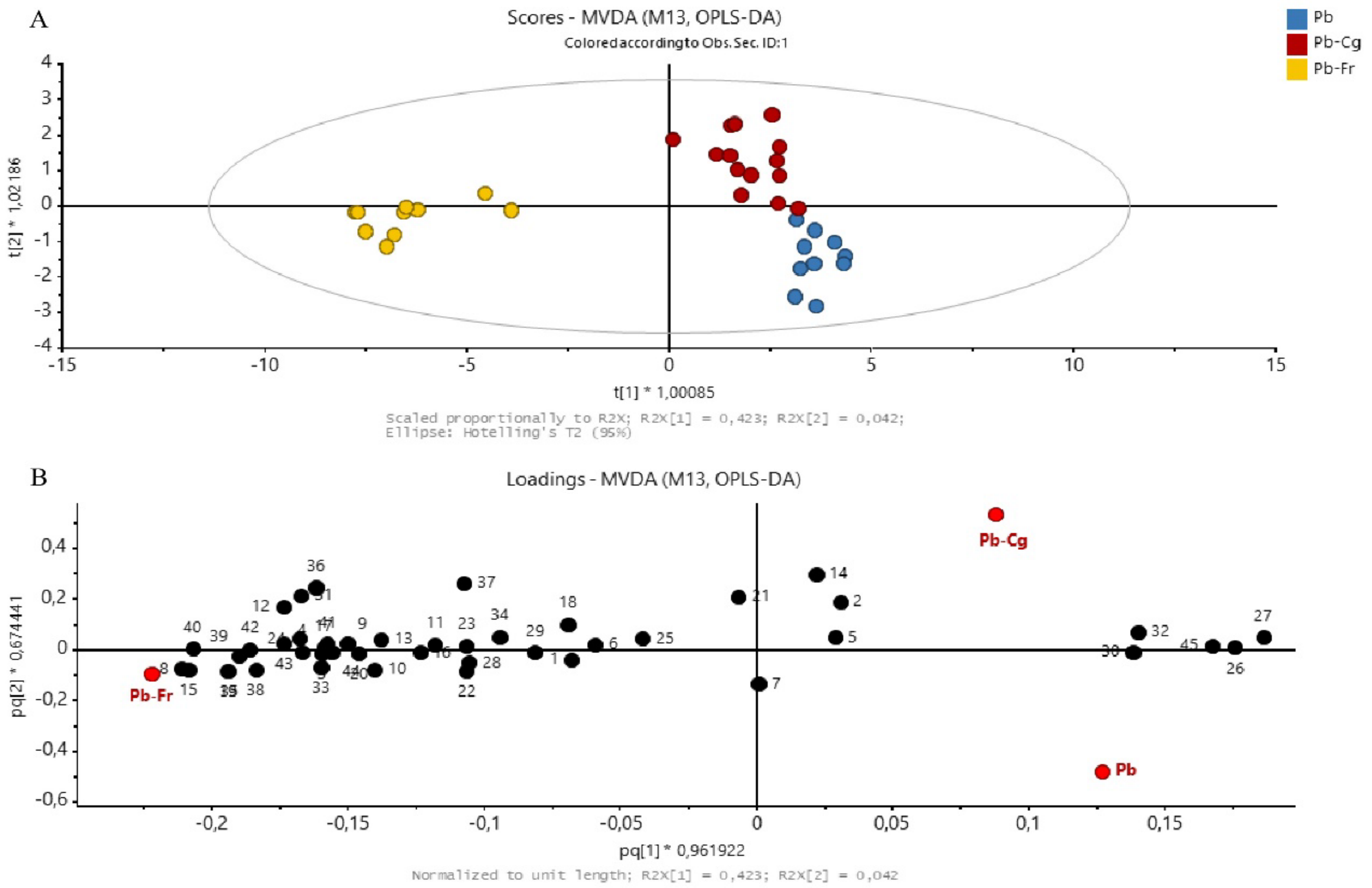


**Fig S9. Overview of volatile blends of *Pieris brassicae* caterpillars, *P. brassicae* caterpillar spit and *P. brassicae* caterpillar frass.** (A), OPLS-DA (Orthogonal Projection to Latent Structures Discriminant Analysis) two-dimensional score plot of treatment groups: Pb, *Pieris brassicae* caterpillars (n = 10); Pb-Fr, *P. brassicae* caterpillar frass (n = 10); and Pb-S, *P. brassicae* spit (n = 12), based on their volatile content. (B), Loading plot showing the contribution of each identified volatile compound to the separation of the different treatments. Volatiles closer to the treatment in the plot indicate higher correlation. Numbers in the loading plot refer to the volatile compounds listed in Supporting information Table S3.

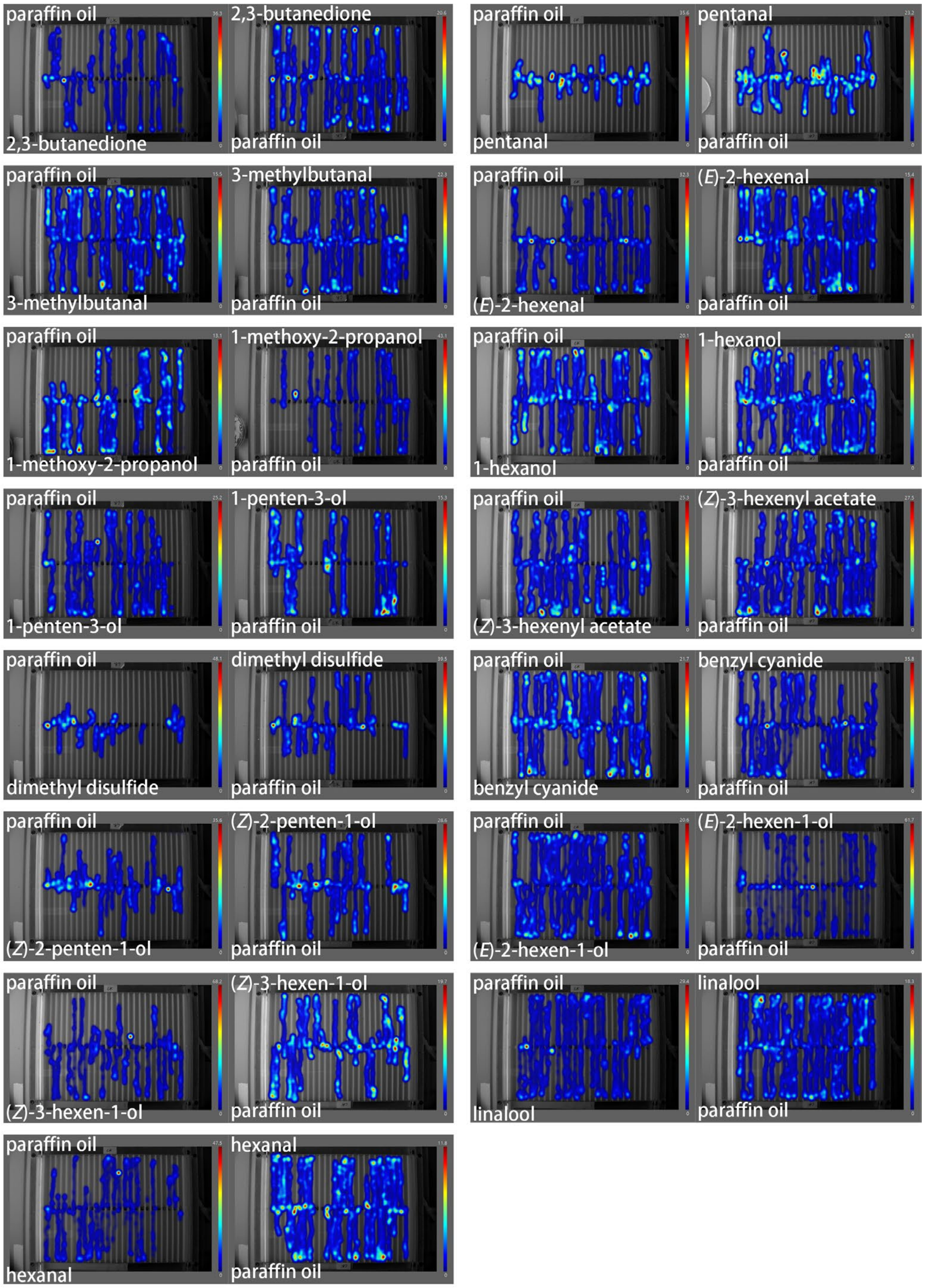


**Fig S10. Heatmaps of WT caterpillar movement in response to the tested chemicals.** The chemical compounds are indicated in the figures and color legend indicates the time (s) that each caterpillar stayed at certain locations.

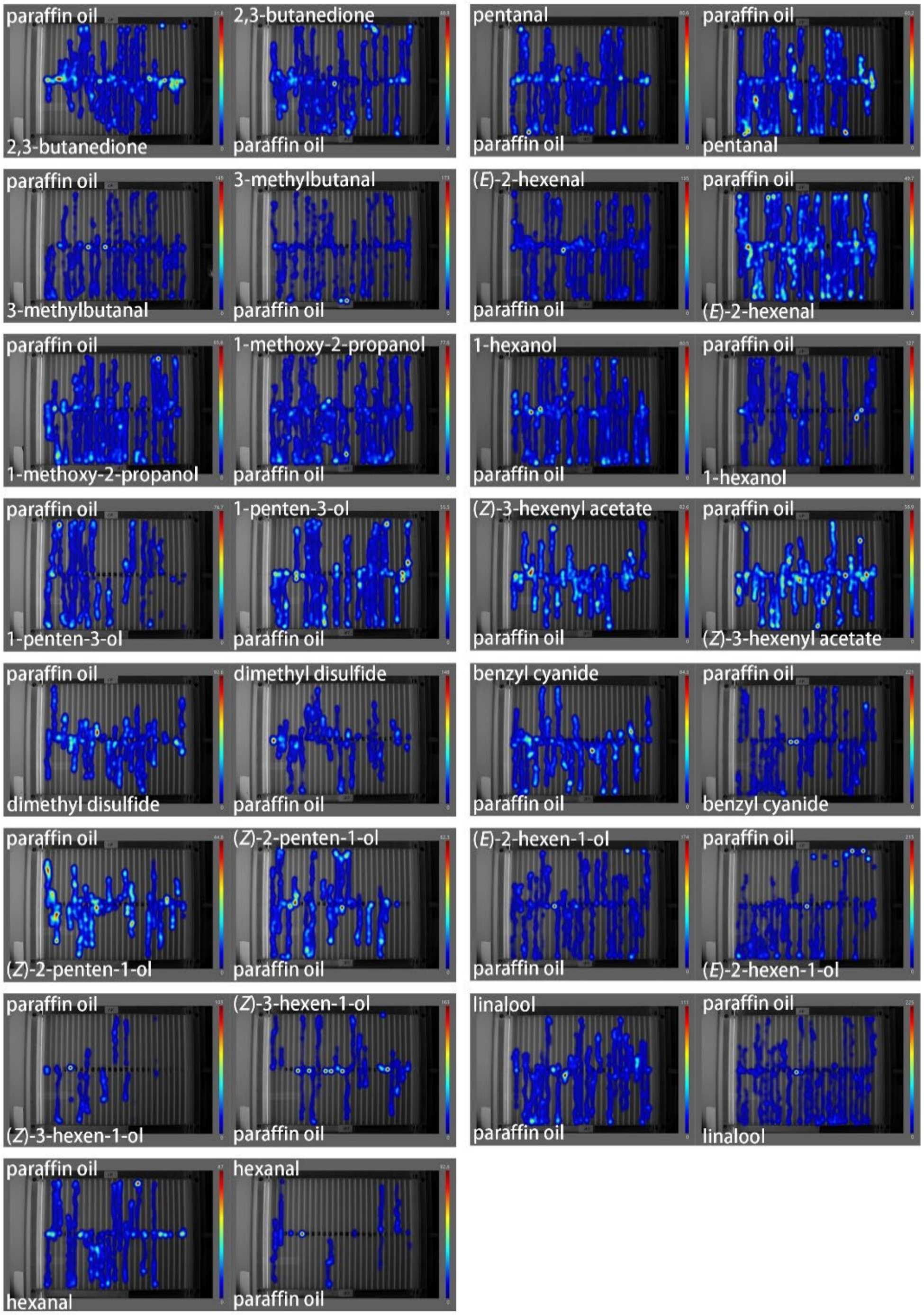


**Fig S11. Heatmaps of *Orco* KO caterpillar movement in response to the tested chemicals.** The chemical compounds are indicated in the figures and color legend indicates the time (s) that each caterpillar stayed at certain locations.
